# Tau isoforms modulate the axon initial segment controlling axonal trafficking and neuronal excitability

**DOI:** 10.64898/2026.09.01.745409

**Authors:** Cayetana Arnaiz, Gonzalo Escalante, Lautaro Rodriguez Donghi, Mariana I Holubiec, Olivia Pedroncini, Indiana Páez Paz, Clara Gaguine, Julieta Bianchelli, Jordi Navarro, Micaela D Garcia, Lucía F Lopez, Alan M Szalai, Antonia Marin-Burgin, Fernando D Stefani, Maria E. Avale, Tomas L Falzone

## Abstract

The axon initial segment (AIS) is a specialized neuronal compartment integrating action potential initiation with selective control of axonal trafficking. The microtubule-associated protein tau is a central regulator of cytoskeletal organization and transport, yet how distinct tau isoforms contribute to AIS development and function remains unclear. Here, we examined the role of tau isoform relative abundance in regulating AIS establishment, maturation, excitability, and transport selectivity using murine primary neurons and human induced pluripotent stem cell (hiPSC)-derived neurons combined with super-resolution imaging, electrophysiology, and live trafficking assays. We found that tau expression levels and isoform content modulate the timing and robustness of AIS maturation. In murine neurons, tau deficiency or predominance of 3-repeat (3R) tau delays Ankyrin-G accumulation and AIS stabilization without preventing AIS formation. hiPSC-derived neurons display an intrinsic AIS developmental program accompanied by progressive changes in tau isoform content. Super-resolution DNA-PAINT reveals that endogenous tau decorates axonal microtubules in discrete nanoclusters with compartment-specific distributions. Modulation of the endogenous 3R/4R tau balance in hiPSC-derived neurons shows that isoform composition, independently of tau levels, regulates AIS positioning and Ankyrin-G organization. Functionally, shifts towards 3R-tau reduce sodium currents, impairs action potential firing, and alters lysosomal transport dynamics within the AIS. Together, these findings identify tau isoform balance as a developmental regulator of AIS maturation, linking cytoskeletal organization to neuronal excitability and transport gating. Because the aberrant alternative tau splicing of exon 10 is a defining feature of primary tauopathies, our results provide mechanistic insight into how imbalanced tau isoforms may contribute to neuronal dysfunction.

**Significant Statement:** The axon initial segment (AIS) plays a central role in brain function by integrating action potential initiation with selective control of axonal trafficking. Disruption of tau splicing, resulting in altered ratios of tau isoforms, is a defining feature of several neurodegenerative diseases, yet its functional consequences remain poorly understood. Here, we show that balanced tau isoform expression is required for proper AIS maturation, neuronal excitability, and transport selectivity. By combining a human neuronal model, super-resolution imaging, electrophysiology, and live transport analysis, our work identifies tau isoform balance as a key regulator of AIS development. These findings provide mechanistic insight into how early tau dysregulation may initiate neuronal dysfunction in tauopathies and highlight the AIS as a potential therapeutic target.

## Introduction

The microtubule-associated protein tau is highly enriched in neurons and essential for cytoskeletal organization, neuronal polarity, and axonal integrity (1). From development to the adult human brain, the content of tau isoforms containing either three or four microtubule-binding repeats is tightly regulated by alternative splicing of exon 10. Imbalances in the 3R/4R tau ratio is a hallmark of neurodegenerative tauopathies such as progressive supranuclear palsy (PSP), corticobasal degeneration (CBD), frontotemporal dementia (FTLD-Tau) and Pick’s disease (PiD), while mixed isoform aggregation occurs in Alzheimer’s disease (AD) (2,3). The axon initial segment (AIS) is a specialized, dynamic cytoskeletal structure regulating electrical activity and axonal transport (4), yet how tau contributes to AIS establishment and function remains poorly understood. Given tau’s ability to modulate microtubule organization and motor accessibility, and that tau isoform imbalance is a defining feature of tauopathies, we hypothesized that early tau isoform composition plays a critical role in regulating AIS structure, transport gating, and electrical output in human neurons, thereby contributing to early neuronal dysfunction in neurodegenerative diseases.

Tau is an intrinsically disordered protein that adopts a structured conformation upon microtubule binding, thereby promoting microtubule assembly, stability, and spatial organization (5–7). In the human adult brain, the *MAPT* gene generates six tau isoforms through alternative splicing, particularly by the inclusion/exclusion of exon 10 leading to either the four (4R) or three (3R) microtubule-binding repeat isoforms. Human neurons predominantly express 3R-tau during early development and progressively reach an approximately equal 3R:4R ratio as they mature, an essential transition for proper neuronal function and brain development (8). The canonical role of tau in microtubule assembly and axonal transport depends on its microtubule-binding affinity, which differs between isoforms (9,10). During neuronal polarization, tau is efficiently sorted into the axon, but it is also detectable in the soma and dendrites (11,12). The tau axonal enrichment is partly maintained by a diffusion barrier at the AIS, which restricts tau retrograde diffusion (13). Isoform-specific microtubule-binding properties and differential post-translational modifications further influence the subcellular localization of tau (14). Although complete tau loss does not overtly disrupt microtubule integrity, subtle defects in neuronal outgrowth have been described, underscoring tau’s regulatory but non-obligatory role in cytoskeletal stability. A physiological role for regulated tau splicing during development has been proposed based on isoform-specific biases in anterograde versus retrograde transport, attributed to 3R- and 4R-tau (9). Disruption of the 3R:4R tau balanced ratio is a defining feature of several neurodegenerative diseases. Accumulation of 4R-tau characterizes FTDP-tau, PSP, and CBD, whereas 3R-tau predominates in PiD and Down syndrome (15). In AD, pathological neurofibrillary tangles may contain 3R- or 4R-tau isoforms (16,17). The pathogenic relevance of tau isoform imbalance is further highlighted by intronic, missense, and silent *MAPT* mutations that alter exon 10 splicing and cause familial tauopathies, stressing that dysregulation of the 3R/4R tau ratio may also contribute to neuronal dysfunction in sporadic tauopathies(18). In addition, tau isoform shifts in human-derived neurons impair the axonal transport of amyloid precursor protein (APP), linking isoform imbalances to pathological APP trafficking and mislocalization (9). Moreover, the overexpression of 3R- or 4R-tau isoforms, or mutations that increase exon 10 inclusion, lead to mitochondrial axonal transport defects (19,20). Together, these observations underscore the importance of tau isoform regulation in neuronal development, polarization, and transport dynamics, raising the question of how isoform imbalance may contribute to early neuronal dysfunction in tauopathies.

The AIS is a specialized proximal axonal domain essential for maintaining neuronal polarity and integrating electrical and transport functions (4,21). Initially described as a region containing tightly bundled microtubule fascicles and electron-dense undercoats beneath the plasma membrane (22), the AIS is now recognized as a highly ordered membrane and cytoskeletal structure. Its assembly is orchestrated by ankyrin-G (AnkG), which anchors voltage-gated sodium (Nav) and potassium (Kv) channels and links them to the underlying βIV spectrin–actin lattice (21). Microtubules within the AIS are organized into parallel fascicles that are stabilized by microtubule-associated proteins promoting axonal specification (23,24). These fascicles are cross-linked by microtubule end-binding proteins (EBs) and AnkG, generating a resilient yet selective cytoskeletal interface (25,26). Functionally, the AIS coordinates action potential initiation through the compartmentalized enrichment of Nav and Kv channels, allowing precise control of electrical firing (27,28). It also acts as a selective filter for vesicular trafficking since dendritic cargos are captured by actin–myosin patches and redirected to the soma, whereas axonal cargos preferentially engage microtubules enriched in specific post-translational modifications that recruit kinesin-1 (29–31). Loss of AIS structural integrity disrupts this transport barrier, leading to somatodendritic proteins invading the axon, impairing both polarity and neuronal outputs (32,33). Indeed, perturbations of AIS scaffolding proteins including AnkG and bIV-spectrin, disrupts ion-channel clustering, AIS integrity, action potential firing, and neuronal excitability (34). Despite being structurally robust, the AIS remains highly plastic. Acute depolarization of hippocampal neurons in culture triggers distal AIS shortening within hours (26,35), whereas chronic changes in network activity reposition the AIS in culture and *in vivo* during development and homeostatic adaptation (36,37). Microtubule dynamics and regulated turnover of ion channels contribute to this activity-dependent remodeling (4). Altered AIS composition or plasticity has been reported in models of epilepsy, AD and other tauopathies, highlighting the AIS as a convergent site of structural vulnerability in disease (38,39). Tau may influence AIS organization, as it promotes the formation of fascicles *in vitro,* reminiscent of those observed at the AIS (40). Its assembly along microtubules modulates motor-based transport in axons, suggesting that tau functions extend beyond passive microtubule stabilization (9,41,42). In primary neuronal cultures, overexpression of tau acetylmimetic mutants destabilizes microtubules within the AIS and leads to tau somatodendritic mislocalization (38). The reduction in hippocampal neuronal firing when mutant P301L tau is overexpressed in mice models was suggested due to a tau-mediated distal relocalization of the AIS (43). Furthermore, tau mutations associated with FTLD-MAPT impair AIS plasticity, resulting in neuronal hyperexcitability (44). Despite these insights, the contribution of 3R/4R tau isoforms to AIS establishment and function remains largely unexplored. Here, we characterized the role of tau isoforms in AIS assembly, transport gating, and neuronal excitability, providing insight into early mechanisms of neuronal dysfunction in tauopathies.

## Results

### Differential Tau content in murine hippocampal neurons affects AIS establishment and maturation

To determine whether tau expression or isoform composition regulate early stages of AIS establishment and maturation, we compared primary hippocampal neurons derived from wild-type (WT) mice, human Tau (hTAU) transgenic mice, and tau knock-out (KO) mice. These models differ in total tau levels and in the relative abundance of 3R- and 4R-tau isoforms. To validate tau expression in these cultures, we analyzed hippocampal neuron homogenates at 14 days *in vitro* (DIV14) by western blot using antibodies against total tau, 3R-tau, and 4R-tau (Fig. 1A). At DIV14, WT neurons expressed tau predominantly as the 4R isoform, whereas hTAU neurons expressed approximately 2.5-fold higher total tau levels than WT neurons, with a predominance of the 3R isoform. Tau-KO neurons lacked detectable tau expression (Fig. 1A, B). This pattern mirrors tau expression profiles reported in adult WT, hTAU, and tau-KO mouse brains (45), Fig. 1C, D). We next asked whether these differences in tau expression and isoform content affected AIS formation by analyzing the axonal accumulation of Ankyrin-G (AnkG) proximal to the soma, using MAP2 staining to define the somatodendritic compartment (Fig. 1E-O). We first examined AIS establishment emerging directly from the soma at DIV5. At this early stage, approximately 25% of WT hippocampal neurons displayed a clearly defined AnkG-positive AIS. In contrast, both hTAU and tau-KO neurons showed a significantly reduced proportion of AIS-positive cells (Fig. 1E, F), indicating a delay in early AIS establishment either when total tau levels or isoform composition is altered. Among neurons that did form an AIS, tau-KO cultures exhibited a significantly increased Start-to-Soma distance compared with WT neurons (Fig. 1E, G), suggesting impaired proximal AIS positioning. In contrast, AIS length at DIV5 was similar across all genotypes (Fig. 1E, H), indicating that tau does not affect initial AIS length formation. We then assessed AIS maturation at DIV14 (Fig. 1I). At this later stage, the proportion of WT neurons exhibiting an AnkG-positive AIS increased to approximately 40%. hTAU and tau-KO neurons reached similar overall proportions, although a significant difference between these two genotypes remained evident (Fig. 1I, J). Notably, both hTAU and tau-KO neurons displayed significantly increased Start-to-Soma distances compared with WT neurons at DIV14 (Fig. 1I, K), consistent with impaired AIS maturation. AIS length remained comparable among genotypes (Fig. 1I, L), suggesting that tau primarily influences AIS positioning and molecular stabilization rather than gross length. In WT neurons, AIS maturation was accompanied by a significant increase in AnkG intensity at DIV14 relative to DIV5 (Fig. 1M). This maturation-associated increase in AnkG intensity was absent in both hTAU and tau-KO neurons (Fig. 1N, O), indicating defective AnkG stabilization. Collectively, these results indicate that both tau expression levels and tau isoform composition contribute to the timing of AIS establishment and to AnkG stabilization during maturation, without being strictly required for AIS formation itself.

**Figure 1:**
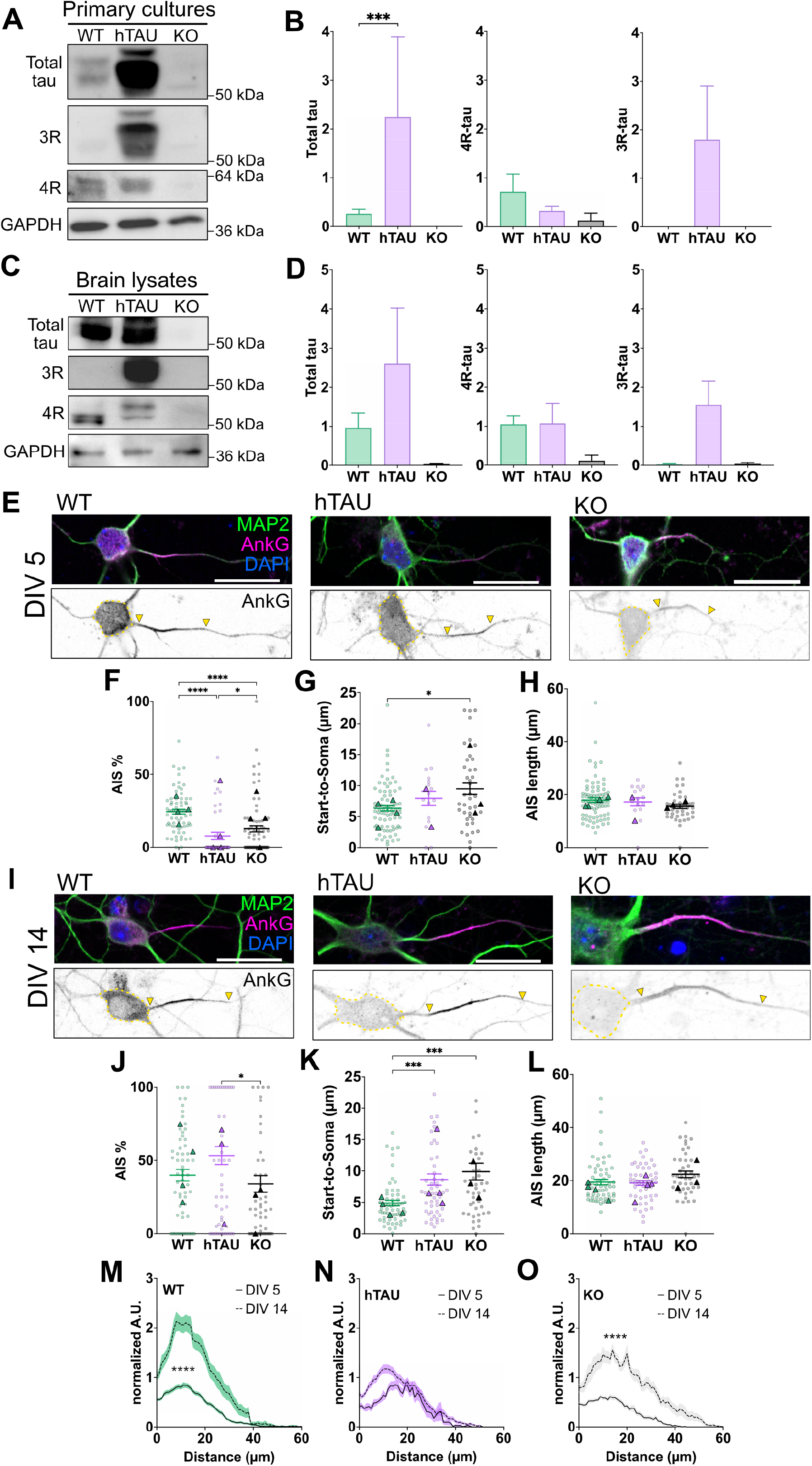
Differential Tau expression in murine hippocampal neurons affects AIS establishment and maturation. Western blots against total tau, and 3R and 4R tau isoforms from (*A*) adult brain cortex and (*C*) hippocampal neuronal culture DIV14 lysates from WT, hTAU, and KO mice. GAPDH was used as a loading control. (*B, D*) Quantification of optical density obtained from the respective western blots representing mean ± SD. Data analyzed by one-way ANOVA (N=2 independent experiments, ***p<0.001). Confocal immunofluorescence images of WT, hTAU, and KO hippocampal neurons at (*E*) DIV5 and (*I*) DIV14, stained with antibodies against AnkG (magenta) and MAP2 (green), to visualize the AIS and the somatodendritic compartment, respectively (scale bars = 20μm). (*F, J*) Quantification of the percentage of neurons with AnkG staining, (*G, K*) the Start-to-soma distance, and (*H, L*) the AIS length in neurons at DIV5 and DIV14, respectively. Kruskal-Wallis tests were applied (N=4, neurons in *F*: n_WT_=69, n_hTAU_=63, n_KO_=73; *G and H*: n_WT_=75, n_hTAU_=14, n_KO_=39; *J*: n_WT_=63, n_hTAU_=56, n_KO_=55; *K and L*: n_WT_=58, n_hTAU_=50, n_KO_=42, from wells obtained from 4 independent experiments, *p<0.05, ***p<0.001, ****p<0.0001). (*M-O*) Plot profile quantification of fluorescence intensity (AU) of AnkG staining at the AIS, for WT, hTAU, and KO neurons at DIV5 and DIV14. Kolmogorov-Smirnov tests were applied. (N=4, neurons in *M*: n_DIV5_=73, n_DIV14_=55; *N*: n_DIV5_=14, n_DIV14_=50; *O*: n_DIV5_=38, n_DIV14_=42, from wells obtained from 4 independent experiments).

### hiPSC-derived neurons exhibit an intrinsic AIS development program

Because tau isoform regulation differs between humans and mice due to species-specific *cis-* and *trans-*acting splicing factors (46), we next examined AIS development in a human neuronal context. To this end, we established experimental settings to study how tau influences AIS formation and maturation within the human genomic and proteomic landscape. We generated forebrain glutamatergic neurons using both pharmacological differentiation and genetic induction protocols, which consistently yielded highly polarized and functionally mature human neurons (Materials and Methods). Across differentiation paradigms, human glutamatergic neurons exhibited a progressive increase in total tau expression over time, accompanied by the progressive increase of both 3R- and 4R-tau isoforms (Fig. 2A, B, Fig. S1). Using Ankyrin-G (AnkG) staining to visualize the AIS, we observed a robust and intrinsic pattern of AIS development. At early stages (DIV5), AIS formation was minimal or absent. In contrast, between DIV25 and DIV37, a substantial proportion of neurons (approximately 60–75%, depending on the protocol) displayed a clearly defined AnkG-positive AIS arising from the soma (Fig. 2C, D; Fig. S1). AIS maturation proceeded gradually over time in culture and was characterized by a progressive reduction in the Start-to-Soma distance, constant AIS length, and a marked increase in AnkG staining intensity between DIV14 and DIV37 (Fig. 2E, F, G, Fig. S1), consistent with progressive AIS stabilization. Importantly, this developmental trajectory was reproducible across differentiation strategies, indicating that AIS establishment and maturation in hiPSC-derived neurons exhibit a reproducible developmental pattern rather than reflecting protocol-specific effects. Together, these observations establish a well-defined temporal window for AIS formation and maturation in human neurons that coincides with the developmental emergence of 3R- and 4R-tau isoforms. This temporal relationship provides a framework for investigating whether changes in tau isoform composition influences AIS structure and function in human neurons.

**Figure 2:**
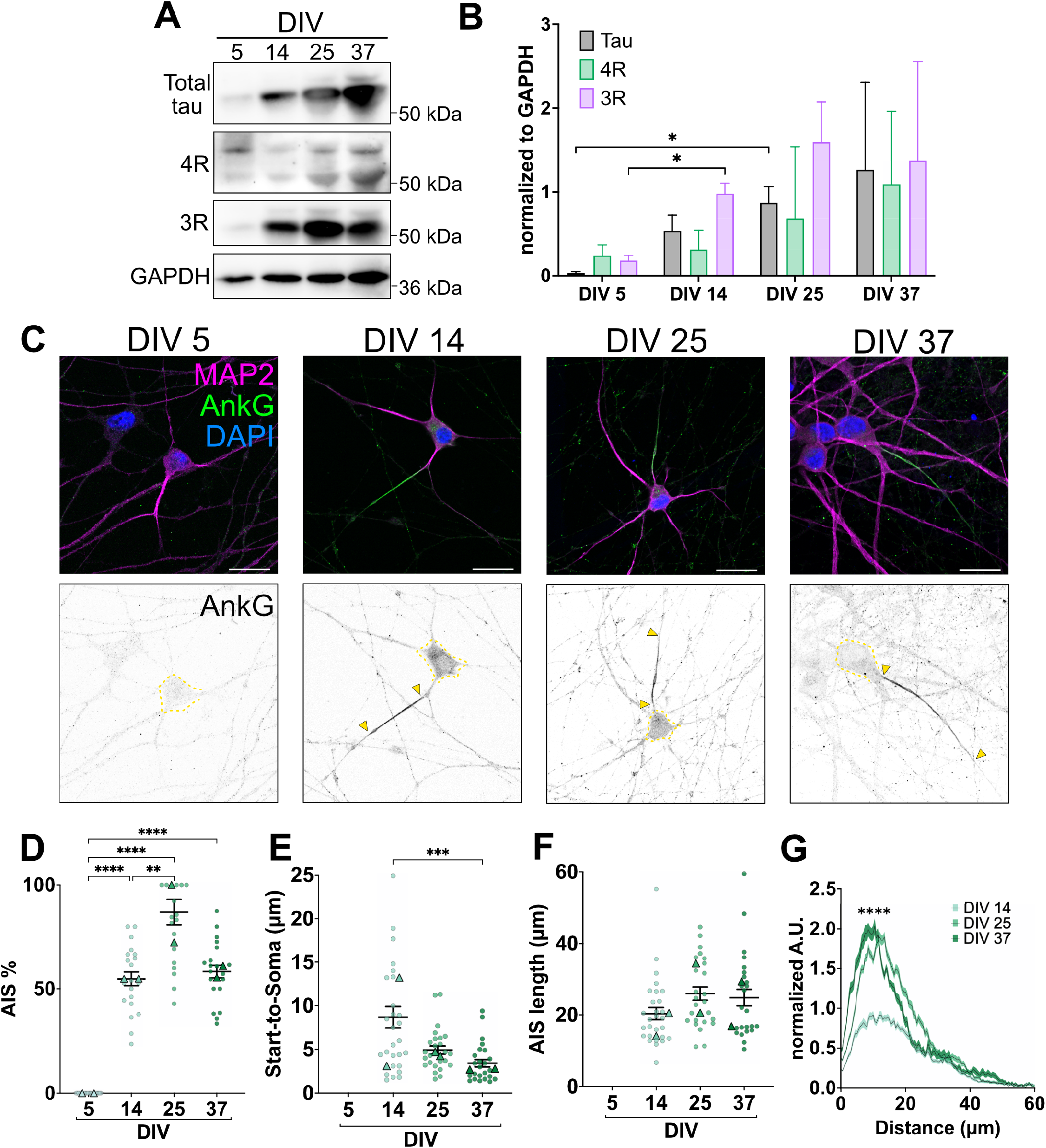
hiPSC-derived neurons exhibit an intrinsic AIS development pattern. (*A-B*) Western blot of i3Neurons homogenates at different time points (DIV5, 14, 25, or 37) showing the expression pattern of total tau, 3R- and 4R-tau isoforms. GAPDH was used as a loading control (N=3 independent experiments, *p<0.05). (*C*) Confocal fluorescence images of i3Neurons at DIV5, 14, 25, and 37 stained against AnkG (green) and MAP2 (magenta), to visualize the AIS and the somatodendritic compartment, respectively (scale bars = 10 μm). (*D*) Quantification of the percentage of neurons with AnkG staining, (*E*) the Start-to-soma distance, and (*F*) the AIS length. Kruskal-Wallis tests were applied (N=2, neurons in *D*: n_DIV5_=20, n_DIV14_=20, n_DIV25_=20, n_DIV37_=20; *E* and *F*: n_DIV5_=0, n_DIV14_=30, n_DIV25_=27, n_DIV37_=26, from wells obtained from 2 independent experiments, **p<0.01, ***p<0.001, ****p<0.0001). (*G*) Quantification of the average arbitrary units of fluorescence intensity (AU) of AnkG staining at the AIS. Kolmogorov-Smirnov test was used (N=2, neurons in *G*: n_DIV5_=0, n_DIV14_=30, n_DIV25_=27, n_DIV37_=26, from wells obtained from 2 independent experiments, ****p<0.0001).

### Differential nanoscale organization of endogenous tau along microtubules in hiPSC-derived neurons

Despite extensive work on tau biology, relatively few studies have examined how tau is spatially organized within neurons, and the nanoscale organization of endogenous, microtubule-bound tau in human neurons remains largely unexplored (47,48). *In vitro* studies have proposed that tau assembles into discrete “islands” or patches along microtubules (49–52), but whether such organization exists in hiPSC-derived neurons in culture is unknown. We therefore sought to characterize the nanoscale distribution of endogenous tau along microtubules in distinct neuronal compartments. To this end, we analyzed hiPSC-derived neurons at DIV14, a stage at which neurons are polarized and AIS formation is underway. An extraction protocol prior to fixation was applied to selectively visualize microtubule-associated tau while minimizing detection of soluble tau (Materials and Methods). At the confocal level, total tau exhibited a broad distribution throughout hiPSC-derived neurons (Fig. 3A). High-magnification imaging of the AIS, identified by AnkG or Neurofascin-186 (NF-186) staining, revealed the presence of total tau as well as both 3R- and 4R-tau isoforms within the AIS region (Fig. 3B–D). To resolve how tau decorates microtubules at the nanoscale level, we next performed 3D DNA-PAINT super-resolution microscopy, which enables single-molecule localization with sub-10-nm precision. We visualized microtubule-bound tau in dendrites, proximal axons encompassing the AIS, and distal axons (Fig. 3E) using the extraction protocol that reduced soluble tau. Super-resolution reconstructions revealed that endogenous tau is organized into discrete nanoclusters along microtubules rather than being uniformly distributed (Fig. 3F). Quantitative analysis showed that tau cluster areas were comparable across dendrites, proximal axons, and distal axons (Fig. 3G, H). In contrast, analysis of nearest-neighbor distances (d1NN) revealed marked compartment-specific differences in cluster spacing. Tau clusters were more widely spaced in dendrites, whereas clusters along axons were more densely packed (Fig. 3I, J). Notably, distal axons exhibited significantly shorter d1NN values than proximal axons, consistent with previously reported proximodistal gradients in tau distribution inferred from lower-resolution studies (11,53). Together, these results demonstrate that endogenous tau adopts a discrete, clustered nanoscale organization along neuronal microtubules in human neurons, with systematic differences in cluster spacing between dendrites and axonal subdomains. This compartment-specific organization provides a structural framework through which tau may differentially modulate microtubule-based transport and AIS function.

**Figure 3:**
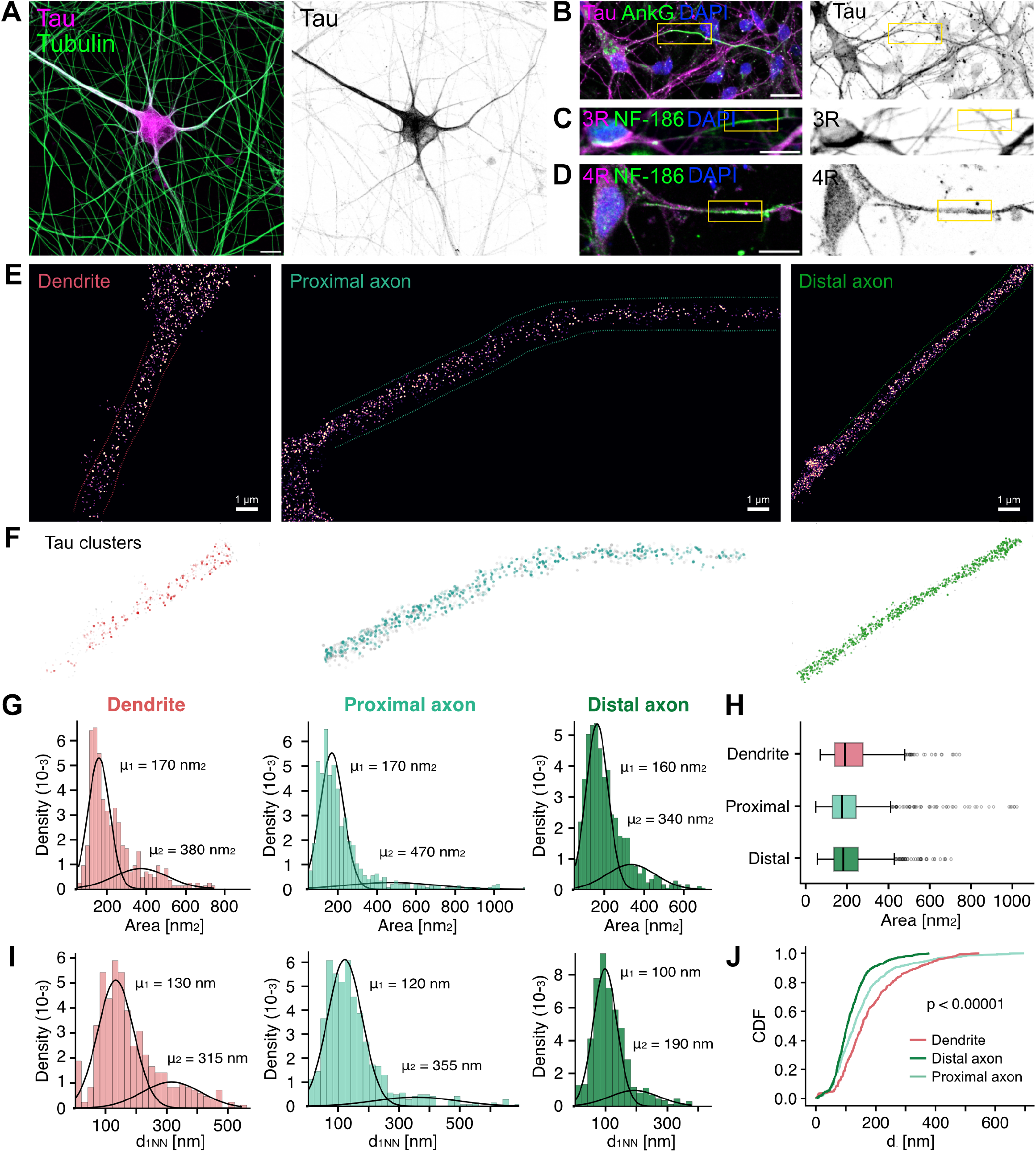
Differential tau nanodistribution along microtubules in neurites of hiPSC-derived neurons. Confocal fluorescence images of i3Neurons stained with antibodies against (*A*) total tau and DM1*α*-tubulin, (*B*) total tau and AnkG, (*C*) 3R-tau and NF-186, and (*D*) 4R-tau and NF-186 (scale bars = 10µm). (*E*) Representative DNA-PAINT rendered images of i3Neurons at DIV14 immunostained against total tau, showing the dendrite, proximal axon and distal axon regions that were used for the analysis. (*F*) Tau clusters identified in dendrite, proximal axon, and distal axon using HDBSCAN and k-means. (*G*) The area of tau clusters and (*I*) the distances to the first nearest neighbor (d1NN), were analyzed in dendrites, proximal axons, and distal axons (n_dendrites_=6, n_prox. axon_=8, n_dist. axon_=6 from 11 ROIs. From 4 independent experiments). In all three regions, frequency histograms were fitted to a two-gaussian mixed model. (*H*) For the comparison of cluster areas, box-plots were graphed, whereas for (*J*) d1NN values, a cumulative frequency analysis (CDF) was performed. For statistical comparisons, Kolmogorov-Smirnov tests were applied (p<0.001). (n_dendrite_ =424, n_proximal_=724, n_distal_=864, from 11 ROIs. From 4 independent experiments).

### Tau isoform composition regulates AIS maintenance and maturation in hiPSC-derived neurons

Results obtained in mouse neurons suggest that tau expression levels and 3R/4R isoform composition influence AIS development. In hiPSC-derived neurons, we observed that tau at the AIS displays a distinct nanoscale organization compared with dendrites and distal axons, and that both 3R- and 4R-tau isoforms are present within the AIS. We therefore asked whether selective modulation of tau isoform composition, without altering tau levels, is sufficient to regulate AIS establishment and maturation in hiPSC-derived neurons. To address this question, we used an established *transplicing* approach that modulates exon 10 inclusion or exclusion in the human *MAPT* primary transcript, thereby shifting endogenous tau contents toward either the 3R or 4R isoform ((9,45,54). hiPSC-derived neurons were transduced at DIV1 with lentiviral vectors LV-PTM3R or LV-PTM4R expressing isoform-specific pre–*transplicing* RNAs, or a control PTM with no *transplicing* ability (Materials and Methods). Western blot analysis of neuronal homogenates at DIV14 confirmed shifts in the 3R/4R tau ratio, without changes in total tau levels when compared with control transduced neurons (Fig. 4A–C). Immunofluorescence staining for AnkG and MAP2 was performed in fixed cultures at DIV14 and 25, to define the AIS and somatodendritic compartments, respectively (Fig. 4D). LV-PTM3R and LV-PTM4R transduced neurons showed similar AIS parameters when analyzed at DIV14 (Fig 4E-H). However, at DIV25 both LV-PTM3R and LV-PTM4R treated neurons showed a reduction in the number of neurons that presented an AIS, suggesting an effect of tau isoform ratio in AIS maintenance (Fig 4E). In addition, while no changes were observed in AIS length at DIV25 we detected a significant difference in AIS Start-to-Soma distance between LV-PTM3R and LV-PTM4R treated neurons (Fig 4F). Moreover, neurons transduced with LV-PTM4R exhibited a reduction in AnkG intensity distribution along the AIS at DIV25 (Fig. 4H, I). Together, these findings show that tau isoform relative content modulates AnkG positioning in hiPSC-derived neurons independently of total tau expression levels, highlighting that 3R/4R tau imbalances affect AIS development and progression.

**Figure 4:**
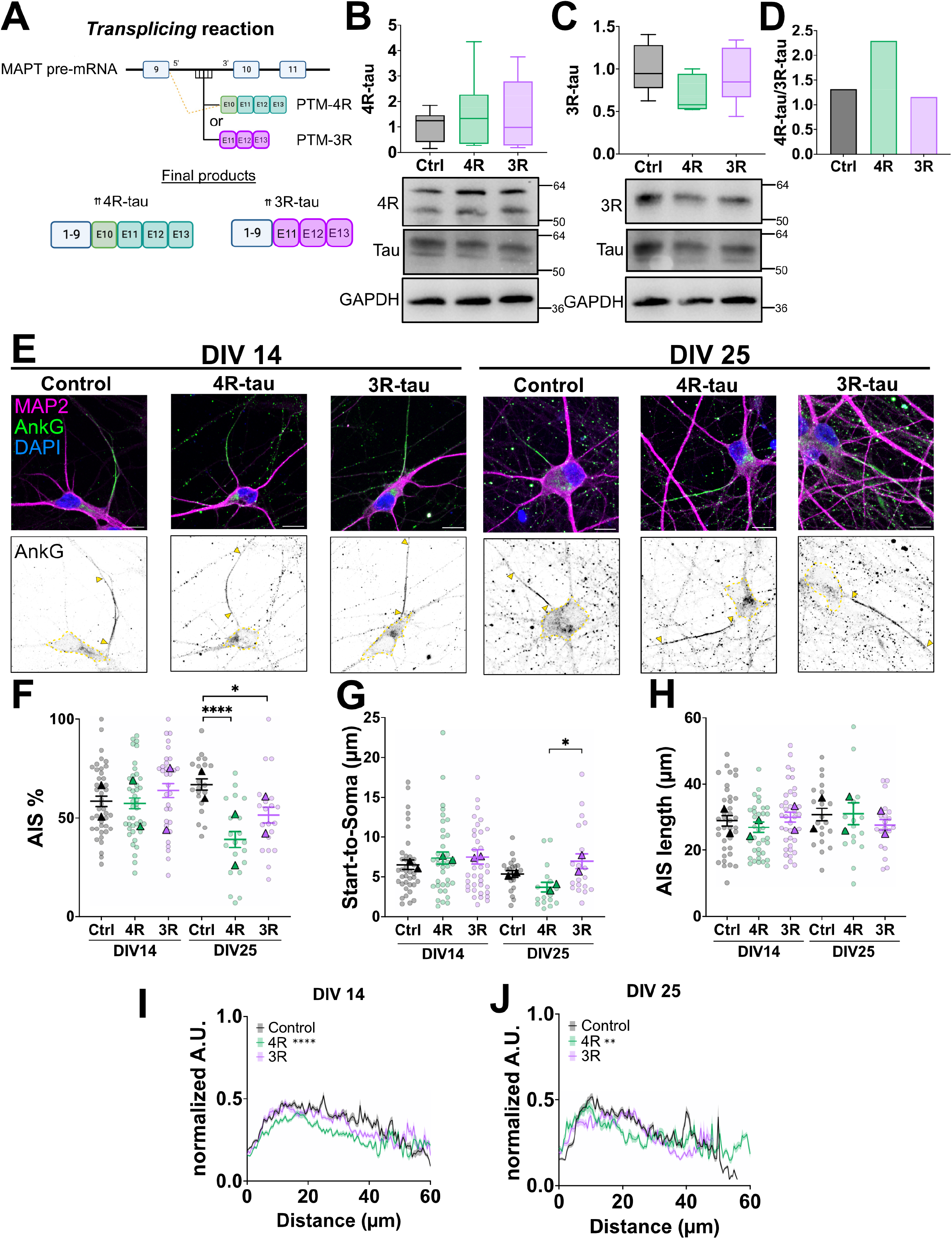
Tau isoforms regulate AIS establishment and maturation in human-derived neurons. (A) Diagram depicting how the *transplicing* system induces the endogenous production of either 4R- or 3R-tau, via transduction of pre-*transplicing* molecules (PTMs) with LV-PTM4R- or LV-PTM3R-tau including/excluding the exon 10 sequence. (*B, C, D*) Western blot of DIV14 i3Neurons homogenates infected with control, LV-PTM4R or LV-PTM3R, showing the expression of 3R- and 4R-tau isoforms. GAPDH was used as a loading control. mean ± SD. Data analyzed by one-way ANOVA (N=6 independent experiments). (E) Confocal fluorescence images of DIV14 and DIV25 i3Neurons infected with control, LV-PTM4R or LV-PTM3R, stained with antibodies against AnkG (green) and MAP2 (magenta) to visualize the AIS and the somatodendritic compartment, respectively (scale bars = 10µm). (F) Quantification of the percentage of neurons with AnkG staining, (G) the Start-to-soma distance, (H) the AIS length, and (I,J) the fluorescence intensity of AnkG staining at the AIS of each neuron. Kruskal-Wallis tests (F*-H*) or Kolmogorov-Smirnov tests (I*, J*) were used (N_DIV14_=6, N_DIV25_=2, neurons in F: n_DIV14_=40, n_DIV25_=20; *G-J*: n_DIV14_=37, n_DIV25_=20, from wells from 2 independent experiments, **p<0.01,***p<0.001,****p<0.0001).

### Tau isoforms composition regulates the electrical properties of hiPSC-derived neurons

Upon observing that tau isoform imbalance had differential effects on the AIS of hiPSC-derived neurons, we tested whether isoform-specific tau changes regulate functional aspects of the AIS, such as the control of electrical activity. We transduced neurons with LV-PTMs at DIV14 and performed whole cell patch-clamp recordings at DIV35 (Fig. 5A–F-). Patch-clamp measurements revealed similar input resistance, action potential amplitudes and threshold values in neurons transduced with LV-PTM3R- or LV-PTM4R-tau when compared with the control condition (Fig. S2). However, neurons transduced with LV-PTM3R-tau exhibited significant reductions in sodium currents relative to controls, while potassium currents remained unchanged (Fig. 5B-E). However, this reduction in sodium conductance was not observed in neurons transduced with LV-PTM4R-tau (Fig. 5B-D). Moreover, the LV-PTM3R-tau condition showed a significant decrease in the number of evoked action potentials compared with both control and LV-PTM4R-tau transduced neurons (Fig. 5F). Since tau influences neuronal polarity and synaptic function, we next examined arborization complexity following tau isoform modulation, by performing Sholl analysis (Fig. 5G). Interestingly, LV-PTM3R-tau significantly reduced neuronal arborization, whereas LV-PTM4R-tau did not show any morphological changes when compared to control (Fig. 5G). Moreover, we evaluated the spine density of the cultures, but neither LV-PTM3R- or LV-PTM4R-tau altered dendritic spine density (Fig. 5H, I), suggesting that the intrinsic changes in neuronal firing mediated by tau isoforms are not due to dendritic structural alterations. Together, these findings indicate that perturbations in tau isoform ratio modulate electrical properties associated with the AIS in hiPSC-derived neurons, with 3R-tau exerting a specific inhibitory effect on sodium currents and action potential generation.

**Figure 5:**
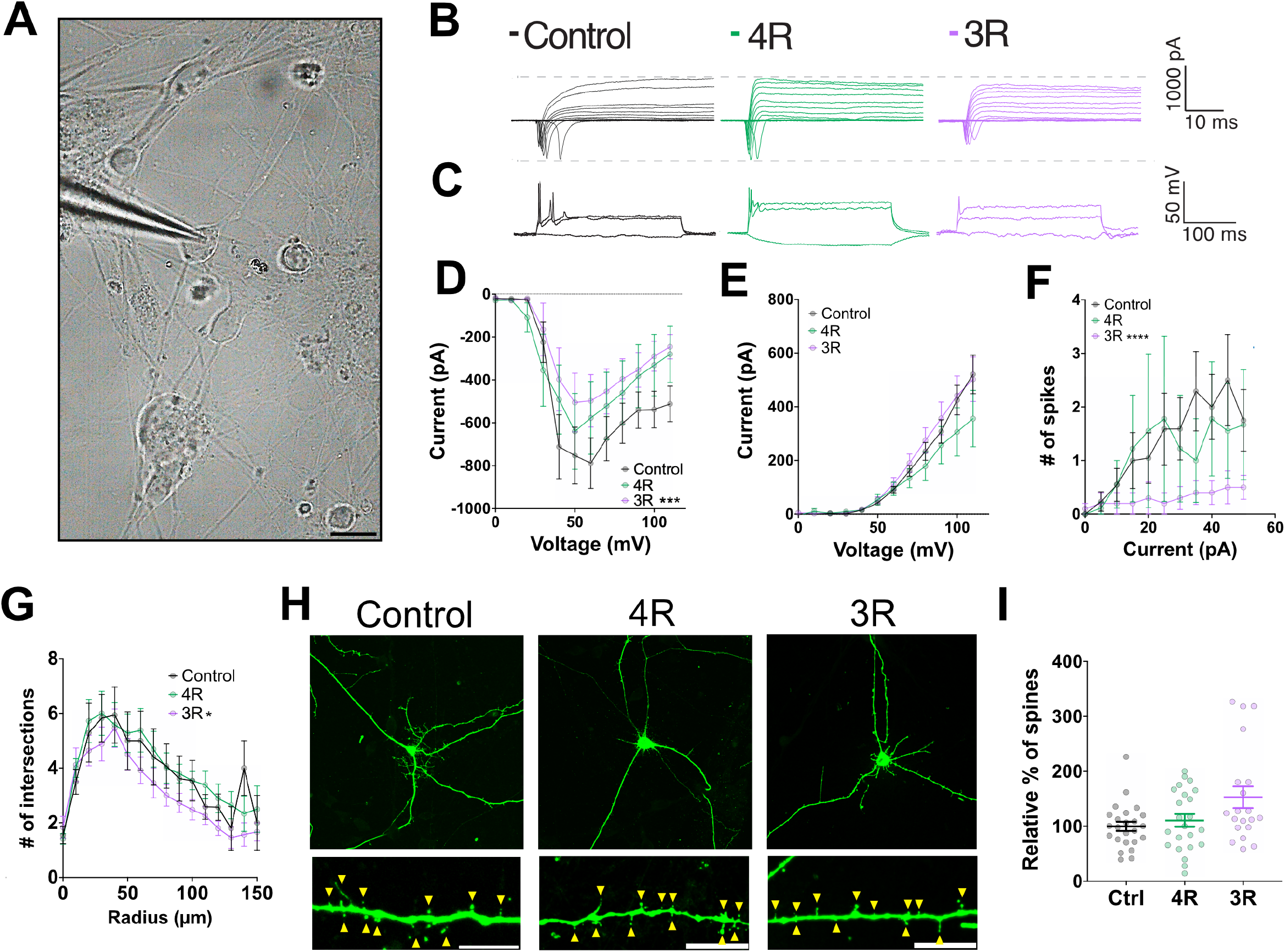
Tau isoforms regulate the electrical properties of hiPSC-derived neurons. (*A*) Brightfield representative image of hiPSC-derived neurons transduced at DIV14 with control, LV-PTM4R or LV-PTM3R (scale bar = 10 µm). Electrophysiological recordings were taken at DIV35 in whole-cell patch-clamp configuration. (*B*) Representative current and (*C*) spike traces obtained after increasing voltage steps. (*D, E*) When performing recordings in voltage-clamp configuration, Na+ and K+ currents were measured for each condition. (*F*) To record the number of evoked firing events upon current injection, current-clamp mode was used. For statistical analysis, ANOVA tests were applied (neurons N_control_=21, N_4R_=9, N_3R_=11, ***p<0.001, ****p<0.0001). (*G*) Sholl analysis to evaluate neuronal arborization. (*H*) Images obtained by confocal microscopy of the entire neuron, and the magnification of a dendrite with spines (scale bars = 10 μm). (*I*) The relative percentage of the number of spines for each condition was obtained. Kruskal-Wallis tests (*G*) or Kolmogorov-Smirnov tests (*I*) were applied (Neurons N_control_=24, N_4R_=23, N_3R_=20, from wells from 3 independent experiments *p<0.05).

### Tau isoforms modulate AIS-dependent barrier and transport functions in hiPSC-derived neurons

Another relevant function of the AIS, linked to its highly specialized structural organization, is the regulation of axonal cargo entry and exit. Consequently, the AIS is considered a selective barrier for the trafficking of axonal organelles and vesicles (35,36,55). To test whether tau isoforms regulate the AIS-associated barrier function, we analyzed lysosomal transport dynamics within the AIS (Fig 6). HiPSC-derived neurons were transduced at DIV1 with LV-PTM3R- or LV-PTM4R-tau, to shift the 3R/4R tau isoform ratio before AIS establishment. Live-imaging experiments were performed at DIV14 under CO_2_ and 37°C controlled conditions. To visualize the AIS in live neurons and simultaneously track lysosomes, we incubated cells with a green fluorescently labeled antibody against the extracellular domain of Neurofascin-186 (liveNF-186) together with LysoTracker-Red prior to imaging (Fig. 6A, B). Thirty-seconds continuous movies were recorded (Movie S1), and lysosomal trajectories were analyzed using ImageJ, Kymotracker, and Kymoanalysis semi-automated programs (58) (Fig. 6C). Neurons transduced with LV-PTM3R-tau exhibited increased total lysosomal density within the AIS compared with control and LV-PTM4R-tau conditions (Fig. 6D). Notably, this increase was driven specifically by a higher number of static lysosomes within the AIS (Fig. 6E). In contrast, the number of pauses in moving vesicles was significantly reduced in both LV-PTM4R- or LV-PTM3R-tau relative to control (Fig. 6F). Lysosomal transport along microtubules relies on the coordinated activity of multiple processive kinesins and dynein motors, described by a “tug-of-war/coordination” model (9,56). Therefore, we quantified anterograde and retrograde segmental velocities of mobile lysosomes. Histograms of segmental velocity distributions were fitted to a three-gaussian mixture model associated with slow (mode A), medium (mode B) and fast (mode C) velocities, as previously established (Fig 6G, Fig. S3). Anterograde segmental velocities were significantly affected by LV-PTM3R-tau, with reductions in the average velocity of the faster modes B and C relative to controls (Fig 6G, Table 1). Retrograde segmental velocities were also significantly reduced in the LV-PTM3R-tau isoform condition, showing reduced average velocity and weight of mode B compared with control (Fig. 6H, Table 1). In contrast, LV-PTM4R-tau only affected the retrograde segmental velocities showing a reduced average velocity in mode A and a decrease in weight of mode B (Fig. 6H, Table 1). Overall, these results suggest that lysosomal transport dynamics within the AIS are modulated by the relative abundance of 3R- and 4R-tau.

**Figure 6:**
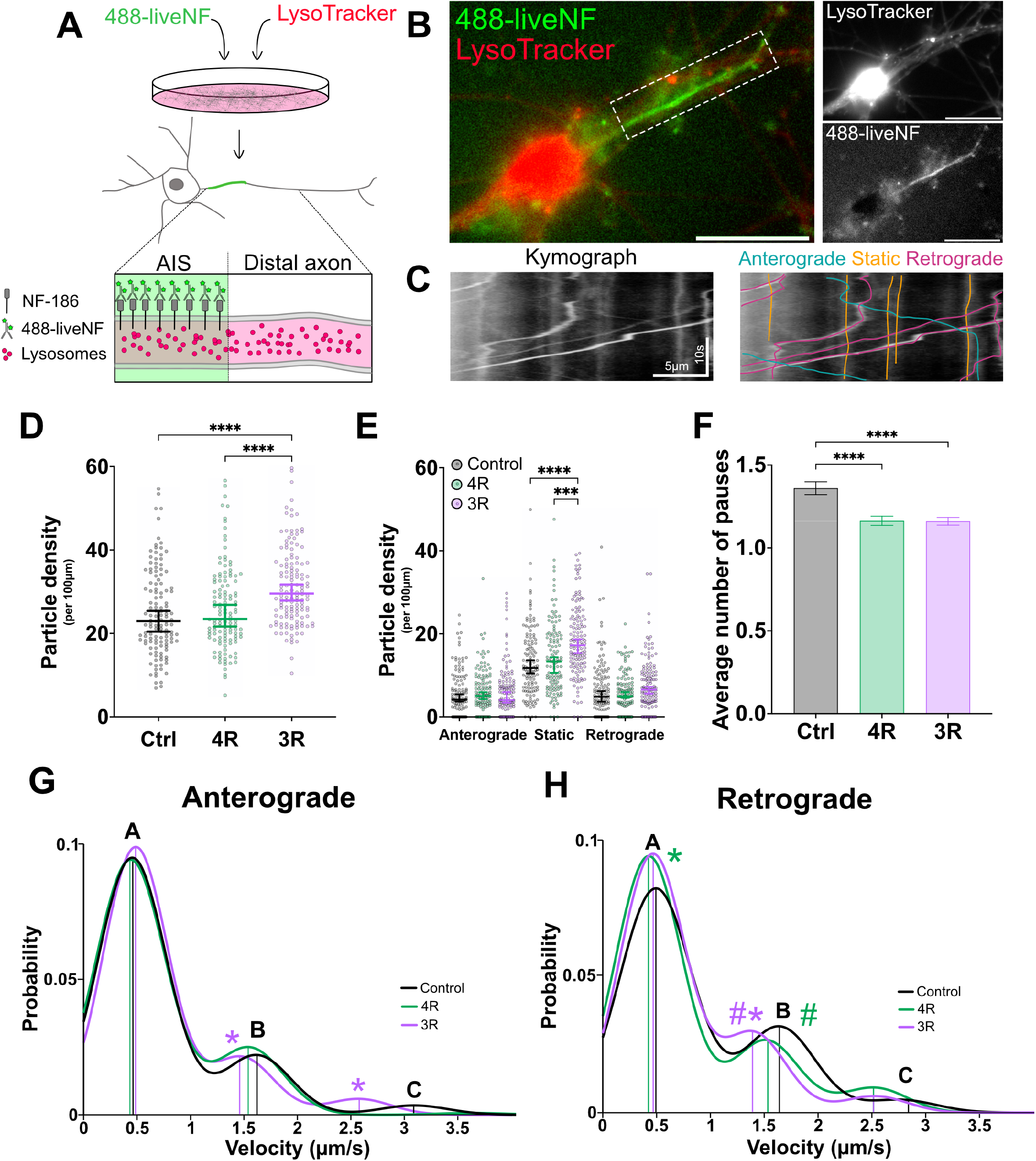
Tau isoforms modulate AIS-dependent barrier and transport functions in hiPSC-derived neurons. (*A*) Glutamatergic neurons were obtained following the SMAD inhibition protocol. At DIV1, the neurons were transduced with control, LV-PTM4R or LV-PTM3R. At DIV14, transduced neurons were treated with LysoTracker to visualize lysosomes, and liveNF-186 was used to localize the AIS. (B) Representative image for a DIV14 neuron showing AnkG (green) and lysosomes (Lysotracker-red) of a continuous 30-second movie. (*C*) Particle movement was analyzed with kymographs obtained in ImageJ, and using the Kymotracker and Kymoanalysis programs. (*D, E*) Particle density and (*F*) the average number of pauses on each condition were obtained. Kruskal-Wallis tests were applied (*D and E*: kymographs N_control_=126, N_4R_=125, N_3R_=136; *F*: particles N_control_=937, N_4R_=838, N_3R_=1228, from wells from 3 independent experiments, ***p<0.001, ****p<0.0001). (*G*) Anterograde and (*H*) retrograde segmental velocities distributions were fitted to a three-gaussian mixed model. Significance was established if the mean velocity µ+/-SD (*), or the weight+/-SD (#) of each mode in the experimental conditions did not overlap with the corresponding modes in the control.

**Table 1.** Anterograde and retrograde segmental velocities modeled as a combination of normal distributions. Values of the anterograde and retrograde central positions and fractions of the mode parameters in control, LV-PTM4R- and LV-PTM3R-tau infected neurons, adjusted to a 3-Gaussian mixed model distribution (A, B, and C). Errors were determined as the SD of n = 1000 resampling with replacement bootstrapping procedure. Fraction mode significance was determined from comparison of the 20% non-overlapping CIs with control condition.

Anterograde
| Mode center |  |  |  |
| --- | --- | --- | --- |
|  | A Max +/- SD | B Max +/- SD | C Max +/- SD |
| Control | 0.455 +/- 0.014 | 1.615 +/- 0.052 | 3.092 +/- 0.188 |
| 4R | 0.430 +/- 0.026 | 1.506 +/- 0.119 | 3.511 +/- 0.662 |
| 3R | 0.483 +/- 0.020 | 1.460 +/- 0.100* | 2.563 +/- 0.145* |
| Mode fraction |  |  |  |
|  | Mode A +/- SD | Mode B +/- SD | Mode C +/- SD |
| Control | 0.788 +/- 0.015 | 0.183 +/- 0.014 | 0.029 +/- 0.006 |
| 4R | 0.777 +/- 0.033 | 0.209 +/- 0.018 | 0.013 +/- 0.021 |
| 3R | 0.780 +/- 0.025 | 0.171 +/- 0.017 | 0.049 +/- 0.014 |

Retrograde
| Mode center |  |  |  |
| --- | --- | --- | --- |
|  | A Max +/- SD | B Max +/- SD | C Max +/- SD |
| Control | 0.490 +/- 0.024 | 1.626 +/- 0.104 | 2.814 +/- 0.320 |
| 4R | 0.427 +/- 0.017* | 1.518 +/- 0.072 | 2.536 +/- 0.102 |
| 3R | 0.463 +/- 0.016 | 1.398 +/- 0.054* | 2.526 +/- 0.114 |
| Mode fraction |  |  |  |
|  | Mode A +/- SD | Mode B +/- SD | Mode C +/- SD |
| Control | 0.692 +/- 0.025 | 0.262 +/- 0.015 | 0.046 +/- 0.025 |
| 4R | 0.725 +/- 0.019 | 0.205 +/- 0.015# | 0.070 +/- 0.014 |
| 3R | 0.727 +/- 0.020 | 0.225 +/- 0.016# | 0.048 +/- 0.009 |

The nanodistribution of tau clusters is significantly denser in distal axons than in the AIS, suggesting that tau may differentially regulate transport in these regions. To assess whether the 3R/4R tau balance exerts distinct effects on lysosomal dynamics outside the AIS, we compared lysosomal density and motility in distal axonal segments and compared them to previous AIS data (Fig. 7A, B). Under control conditions, lysosomal density was significantly lower in distal axons than in the AIS (Fig 7C), reinforcing the concepts of the AIS acting as a selective transport barrier (55). Strikingly, these AIS-distal axonal transport differences were not observed in both LV-PTM3R and LV-PTM4R transduced neurons (Fig. 7C). Furthermore, in control neurons, anterograde transport velocities were slower in distal axons compared with the AIS, a difference only maintained in LV-PTM3R-tau condition (Fig. 7D). Comparisons between LV-PTM3R and LV-PTM4R transduced neurons revealed that 3R-tau drives faster anterograde velocities and slower retrograde velocities (Fig. 7E). This pattern agrees with previously described effects on axonal APP vesicle transport when 3R/4R tau balance is perturbed (9). Together, these findings demonstrate that tau isoforms exert differential effects on lysosomal transport at the AIS and at distal axons, in a manner that correlates with the compartment-specific nanodistribution of tau clusters.

**Figure 7:**
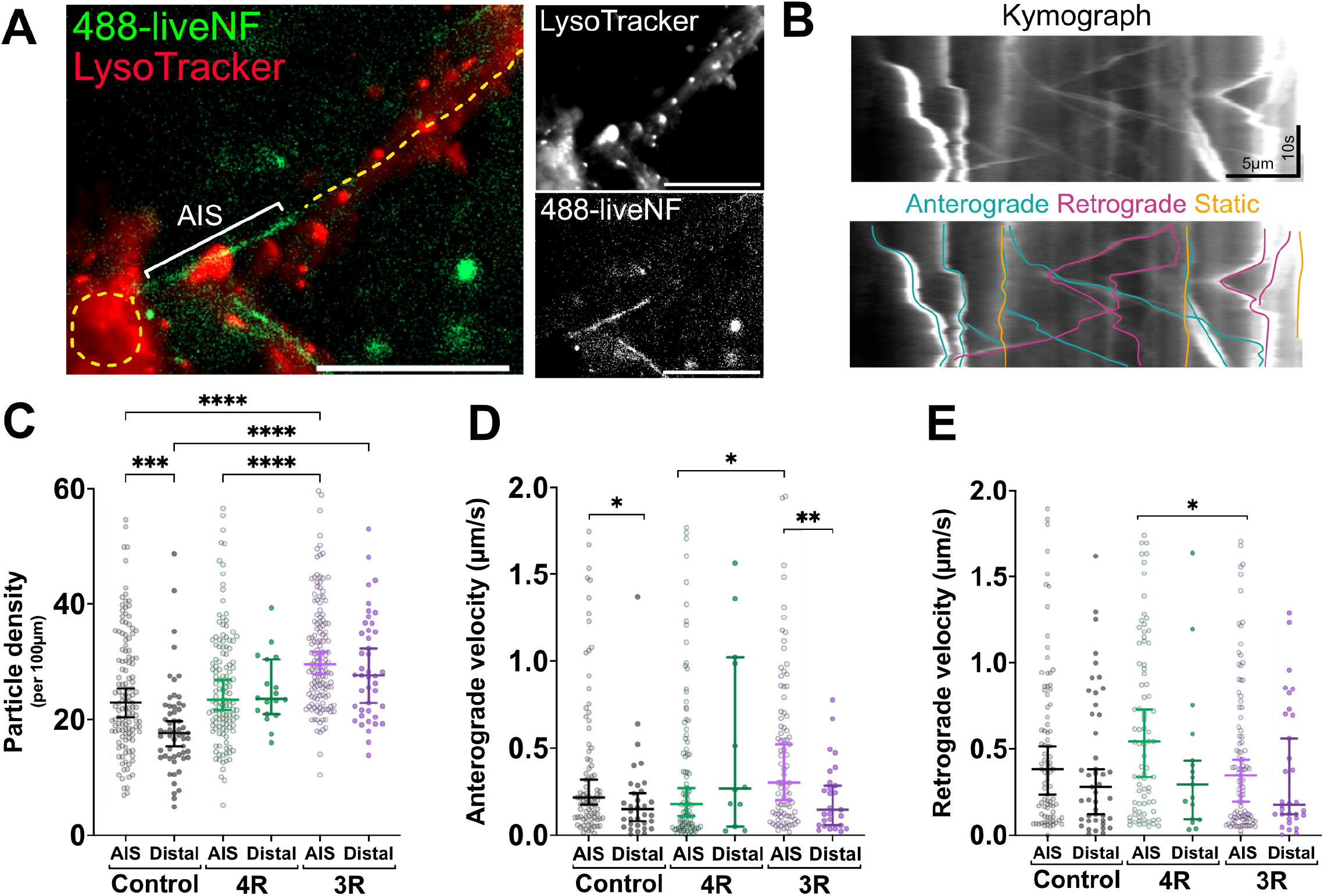
Tau isoform content differentially affects lysosomal transport outside the AIS. (*A*) Glutamatergic neurons were obtained following the SMAD inhibition protocol. At DIV1, the neurons were transduced with control, LV-PTM4R or LV-PTM3R. At DIV14, the transduced neurons were treated with LysoTracker to visualize lysosomes, and liveNF-186 was used to localize the AIS. Representative image for a DIV14 neuron showing AnkG (green) and lysosomes (Lysotracker-red) of a continuous 30-second movie. (*B*) Particle movement in the region distal to liveNF-186 staining was analyzed with kymographs obtained in ImageJ, and using the Kymotracker and Kymoanalysis programs. (*C*) Particle density, (*D*) anterograde velocities, and (*E*) retrograde velocities were compared between lysosomes in the AIS and outside the AIS (termed as “Distal”). Kruskal-Wallis tests were applied (*C*: Kymographs N_control_=57, N_4R_=18, N_3R_=43; *D*: Kymographs N_control_=34, N_4R_=12, N_3R_=27; *E*: Kymographs N_control_=45, N_4R_=17, N_3R_=32, from wells from 3 independent experiments, *p<0.05, **p<0.01,***p<0.001,****p<0.0001).

## Discussion

The AIS is a critical integrator of neuronal polarity, excitability, and compartment-specific trafficking (4). While tau dysfunction has long been implicated in altered microtubule dynamics, neuronal firing, and intracellular transport, how developmentally regulated tau isoform balance influences AIS maturation and function has remained unclear. In this study, by combining murine and human neuronal cultured models with super-resolution imaging, electrophysiology, and live transport assays, we identify tau isoform composition as a regulator of AIS maturation, linking cytoskeletal organization to neuronal excitability and selective transport gating.

Tau isoform expression, particularly the balance between 3R- and 4R-tau, is dynamically regulated during development but becomes disrupted in tauopathies such as PSP, CBD, and AD (2,57). Tau dysfunction, through isoform imbalance, post-translational modifications, or mutations, has been shown to affect microtubule stability, neuronal excitability, and intracellular trafficking (38,43,44,58,59). However, the molecular pathways linking tau isoforms to AIS maintenance and function have remained poorly defined.

Our results in murine hippocampal neurons support a model in which tau isoforms modulate the timing and stabilization, rather than the absolute formation of the AIS. Neurons lacking tau or overexpressing predominantly 3R-tau exhibited delayed AIS establishment, altered AIS positioning, and impaired accumulation of Ankyrin-G (AnkG), particularly at later developmental stages. Importantly, the AIS is eventually formed in both hTAU and tau-KO neurons, indicating that tau is not strictly required for AIS assembly. This finding is consistent with the presence of redundant microtubule-associated proteins such as TRIM46, DCX, and MAP7, which are sufficient to support initial microtubule fasciculation and AIS formation (23,24). Instead, tau isoforms appear to fine-tune AIS maturation by promoting microtubule bundle stability and AnkG enrichment during a defined developmental window.

Given interspecies differences in *MAPT* splicing regulation (60), we extended our analysis to hiPSC-derived neurons. HiPSC-derived neurons exhibited a conserved AIS maturation trajectory characterized by progressive increases in the proportion of AIS-positive cells, stabilization of AIS length, reduction in the Start-to-Soma distance, and increased AnkG intensity, in agreement with prior developmental studies (61). We detected 4R-tau as early as DIV14, and increases as time progressed, likely due to the NGN2-induced differentiation protocol (62,63).

A central advance of this study is the characterization of the nanoscale organization of endogenous, microtubule-bound tau in human neurons. Using 3D DNA-PAINT super-resolution microscopy, we show that tau is organized into discrete nanoscale clusters, with inter-cluster spacing decreasing along the axon. This organization extends and refines the classical view of tau as a uniformly distributed microtubule-associated protein (53,64), and differs from tau “islands” described in *in vitro* systems (49,50,52). Prior super-resolution studies largely relied on tau overexpression and focused on oligomers or aggregates (47,65), whereas our analysis reveals the native nanoscale arrangement of endogenous tau along microtubules in hiPSC-derived neurons. The compartment-specific spacing of tau clusters, combined with conserved internal cluster organization, suggests intrinsic spatial regulation that may underlie regional differences in motor accessibility and transport modulation.

By using the *transplicing* system we shifted 3R/4R tau content in hiPSC-derived neurons without overexpression artifacts (9). Both LV-PTM3R-tau and LV-PTM4R-tau conditions impaired the AIS presence at DIV25, confirming a developmental requirement for tau isoform content regulation. In addition, LV-PTM3R and LV-PTM4R transductions induced differential positioning of the AIS that could be associated with its structural integrity and electrical function.

Functionally, tau isoform imbalance modulates AIS-dependent neuronal excitability. LV-PTM3R -tau selectively reduced sodium currents and action potential firing without affecting passive membrane properties or dendritic spine density, pointing to a primary AIS defect. These findings are consistent with prior evidence that tau mutations, depletion, hyperphosphorylation, or extracellular tau species alter AIS structrure, plasticity and disrupt Nav channel anchoring (43,44,66), likely through impaired AnkG organization (27,28,67). Thus, tau may modulate excitability through two non-mutually exclusive mechanisms: directly, by perturbing the AnkG-associated AIS scaffold and Nav-channel organization; and/or indirectly, by altering synaptic input through tau–Fyn regulation of GluN2B-containing NMDA receptors, thereby regulating drive and network excitability (68).

Tau isoforms also modulated the selective transport barrier function of the AIS. LV-PTM3R-tau increased lysosomal density within the AIS, particularly in stationary vesicles, and altered both anterograde and retrograde transport dynamics, whereas LV-PTM4R-tau did not change the density of lysosomes at the AIS but selectively impaired the retrograde segmental velocities. Moreover, the reduction in the number of pauses observed for both tau isoform shifted conditions suggests an alteration in the AIS barrier function. This alteration is also supported by the fact that isoform imbalance abolished physiological differences in transport behavior between the AIS and distal axon compartments, consistent with a loss of AIS-specific gating (55). The distinct effects of LV-PTM3R- and LV-PTM4R-tau on lysosomal transport contrast with previously described tau isoform-dependent modulation of APP vesicle trafficking in distal axons (9), suggesting compartment-specific motor coordination mechanisms, potentially reflecting differential dynein sensitivity to tau-decorated microtubules (50).

Together, our findings support a model in which balanced tau isoform content acts as a developmental regulator of AIS maintenance, coordinating cytoskeletal organization, excitability, and selective transport. A shift towards 3R-tau produces a functional state consistent with an immature AIS, while deviations toward 4R-tau also perturb neuronal homeostasis, underscoring the importance of regulated isoform balance rather than a single “pathogenic” isoform. Because disrupted tau splicing is a defining feature of primary tauopathies (2,57), these results provide mechanistic insight into how early isoform imbalances may initiate neuronal dysfunction. More broadly, this work identifies the AIS as a critical site through which tau isoforms influence neuronal physiology and suggests AIS-centered pathways as potential targets for therapeutic intervention in tau-mediated neurodegeneration.

## Materials & Methods

### Antibodies

Monoclonal antibodies against the following proteins were used: AnkG (mouse, clones N106/36, NeuroMab, 1:100), *α* -tubulin (mouse, clone DM1*α*, Sigma T9026, 1:500), 3R Tau (mouse, clone 8E6/C11, Millipore, 1:50 or 1:100), 4R Tau (mouse, clone 1E1/A6, Millipore, 1:50 or 1:100), GAPDH (mouse, AB8245, Abcam, 1:10,000). Polyclonal antibodies against the following proteins were used: MAP2 (rabbit, sc-20172, Santa Cruz Technologies, 1:300), Tau (rabbit, A0024, Dako, 1:300), Neurofascin-186 (rabbit, AIP-025, Alomone Labs, 1:100). The following secondary antibodies were used: rabbit Alexa Fluor 568 (Invitrogen, A11011, 1:500), rabbit Alexa Fluor 488 (Invitrogen, A11008, 1:500), mouse Alexa Fluor 568 (Invitrogen, A11004, 1:500), mouse Alexa Fluor 488 (Invitrogen, A11001, 1:500), HRP goat anti-rabbit (Vector Laboratories, 111-035-144, 1:3000), HRP goat anti-mouse (Vector Laboratories, 115-035-146, 1:3000).

### Mice and primary neuronal cultures

Htau transgenic mice, in a C57BL/6 background, were obtained from Jackson Laboratories (Bar Harbor, Maine, United States; B6.Cg-Mapttm1 (EGFP) Klt Tg (*MAPT*)8cPdav/J. Stock number: 005491). This strain carries a full-length human MAPT BAC transgene on an endogenous mouse Mapt knockout background. Mice were genotyped by PCR to confirm the presence of the human MAPT transgene in heterocigous and the mouse Mapt null background, as previously described (47). The strain was subsequently maintained in our colony with periodic backcrossing to C57BL/6 mice. For the generation of primary neuronal cultures, heterozygous hTau mice were intercrossed to generate sibling offspring carrying the hTau transgene or the endogenous Mapt null allele. Thus, hTau and tau-KO neurons were obtained from littermates generated from the same breeding scheme and were processed in parallel under identical culture conditions. C57BL/6 wild-type (WT) mice carrying the endogenous mouse Mapt allele were obtained from littermates generated during the C57BL/6 backcrossing of the colony and were used as the WT control group. This breeding strategy ensured that the neuronal cultures used for comparisons were derived from closely matched genetic backgrounds and, where applicable, from siblings from the same crosses. The origin and breeding strategy of these mouse lines are consistent with those used in our previous studies (45,66,69). Newborn hippocampal brain regions from WT and hTau mice crosses were dissected on post-natal day 1. Treated independently, hippocampi were incubated in 0.22μm-filtered mixture of 45U papain (Worthington) in PBS, DL-cysteine (Sigma), BSA and glucose (Sigma) enriched with 0.05% of DNase (Boehringer Mannheim) for 30 min at 37 oC. Hippocampi were triturated by carefully pipetting in 10% FBS/DMEM. Cells were grown in 500 uM L-glutamine and neurobasal media supplemented with B27 (Invitrogen) over poly-D-lysine coated coverslips (70).

### hiPSC culture and maintenance

Human induced pluripotent stem cells (hiPSCs) were maintained either on inactivated mouse embryonic fibroblast (MEF) feeder layers in KO-DMEM supplemented with KnockOut Serum Replacement, MEM-NEAA, GlutaMAX, Pen-Strep, β-mercaptoethanol, and bFGF, or under feeder-free conditions on Matrigel in StemFlex medium. Cells were passaged using trypsin or Accutase and plated in the presence of ROCK inhibitor (Y-27632), which was removed after 24 h. hiPSCs expressing doxycycline-inducible neurogenin-2 (NGN2-hiPSCs) (65) were kindly gifted by Dr. Juan Bonifacino. NGN2-hiPSCs were cultured as previously described (71).

### Neuronal differentiation

Cortical forebrain neurons were generated via embryoid body (EB) formation followed by neural induction and neural rosette formation. Rosettes were isolated, dissociated, and plated on poly-ornithine/laminin-coated coverslips in neural differentiation medium supplemented with neurotrophic factors. Glutamatergic neurons were derived using dual SMAD inhibition and cyclopamine during EB induction, followed by rosette formation and neuronal maturation under similar conditions. NGN2-hiPSCs were differentiated into i3Neurons by doxycycline-induced NGN2 expression and cultured in BrainPhys-based neuronal medium supplemented with neurotrophic factors (63).

### Immunocytochemistry and microscopy

For axon initial segment staining, neurons were fixed at different DIV with 4% paraformaldehyde (pH 7.4) in 3% sucrose, 60 mM PIPES, 25 mM HEPES, 5 mM EGTA, and 1 mM MgCl_2_, for 20 min at RT. Permeabilization was carried out in 0.25% Triton X-100 in PBS for 5 min at RT and the block step was carried out in 10% Normal Goat Serum (NGS) in PBS for 1 hour at RT. Primary antibody incubation was done in 1% NGS and 0.1% Triton X-100 in PBS, overnight at 4°C, then three 5 min washes with PBS 1X were done, after which coverslips were incubated with the appropriate Alexa Fluor secondary antibodies for 2 hours at RT, diluted in PBS 1X. Images were taken in a Zeiss LSM710 confocal microscope, using a Plan-Apochromat 40X/1.4 oil DIC M27 objective or a Zeiss epifluorescence microscope, using the same objective, and an Axiocam HRm-3, for AIS presence quantification. The lasers used for confocal imaging were a diode 405-30 for DAPI, an argon laser for Alexa488, and a HeNe543 laser for Alexa 568. Images were 1504 by 1504 pixels, with a digital pixel size of 70.66 nm. For z-stacks, the step size was 360.98 nm.For cytoskeletal staining, neurons were first incubated with an extraction buffer (0.3% Triton X-100 and 0.1% glutaraldehyde in BRB80 buffer) for 1.5 min at RT. Following extraction, a fixation solution (4% paraformaldehyde in BRB80 buffer) was applied for 20 min at RT. After fixation, samples were incubated in permeabilization buffer (0.25% Triton X-100 in BRB80 buffer) for 10 min at RT. BRB80 buffer (pH 6.8) consisted of 80 mM PIPES, 1 mM EGTA, and 4 mM MgCl₂. All subsequent steps, including antibody incubations and washes, were performed as described previously.

### Axon initial segment analysis

AIS analysis was performed in ImageJ by first tracing a segmented line along the axon containing AnkG staining, starting at the soma and then continuing until AnkG staining was indistinguishable from background. Using the intensity profile of the AnkG, Start-to-Soma parameters were delimited, as the distance from the soma to the point at which the change in AnkG intensity was ∼6 times higher than the background (i.e. if we had a 5 A.U. background intensity, the point at which the signal became 30 A.U. was considered the start of the AIS). In the case of the end of the segment, a ∼2 to 3 times decrease of the AnkG intensity was considered as the end point of the AIS. This smaller difference was taken into account since AnkG staining did not abruptly disappear along the axon. Only AIS coming from cell body were accounted in our quahntification and those arising from primary dendrites were discarded. The AIS length was calculated as the difference between these two distances. Finally, AnkG staining intensity was quantified using the plot profile tool by drawing a line along the axon, including the AnkG-positive region up to the soma. AnkG intensity values were normalized to the average intensity measured within each experiment. All image analysis was performed blindly, where each image was numerically named during acquisition and after analysis, each condition was traced back.

### Viral transductio

Neurons were transduced one day after plating (DIV1) for approximately 16hrs with ∼200µl of media, as previously reported (9). Briefly, lentiviral particles containing PTM4R, PTM3R or a control PTM were prepared as previously described (9). PTM3R lacks exon 10 driving 3R-tau isoform synthesis, while PTM4R contains exon 10, leading to 4R-tau protein production. Control PTM lacks a binding domain to the endogenous *MAPT* transcript, so it does not allow trans-splicing (9, 47, 56). PTMs are expressed under the human synapsin promoter in a replication-defective lentiviral vector (LV) prepared as described previously (45). An LV particle expressing GFP was also used as a transduction efficiency control. After 16 hours, the cells were topped up with 300 µl of fresh neuronal medium. 13 to 24 days after LV transduction, neurons were either fixed for immunostaining, lysed for western blotting, or used for live-cell imaging experiments.

### Live-cell imaging of lysosomal trafficking at the AIS

Imaging of live cells and kymograph generation were performed as previously described (72). Briefly, 30 s movies of LysoTracker Red DND-999 (Sigma) moving lysosomes in neurons were recorded using an inverted epifluorescence microscope (Zeiss) connected to an Axiocam HRm-3. Cultures were observed under a 63X/NA1.4 objective and maintained at 37°C, 5% CO_2_, and 10% humidity using a CO_2_ humid chamber and heated stage (Zeiss). To analyze lysosomal trafficking strictly at the AIS, neurons were incubated with an antibody against the extracellular domain of Neurofascin-186 (liveNF-186) labelled with AlexaFluor 488 (a kind gift from Dr. Juan Bonifacino (NIH)), for 30min. Kymographs were generated from the recordings with ImageJ using the multiple kymograph plug-in, and lysosomal dynamics was analyzed using Kymoprocessor, a semiautomated tracking toolbox developed by our group (56).

### Dendritic spine analysis

Neurons were transfected at DIV18 with a plasmid expressing GFP under the CAG promoter (pCAG-GFP) to be able to visualize the whole neuronal cytoplasm and therefore observe dendritic spines. Transfections were performed using 2 µg of plasmid DNA and 2.5 µl of Lipofectamine 2000 (Invitrogen), diluted in OptiMEM (Invitrogen) as the transfection medium. At DIV25, cells were fixed and dendritic spines were imaged using a Zeiss LSM710 confocal microscope, with a Plan-Apochromat 40X/1.4 oil DIC M27 objective. Quantification was performed blindly, by manually counting the number of spines along a 70-100 µm dendritic segment, normalizing the number of spines to the length of the analyzed dendritic segment. Spine density changes were expressed as percentage change relative to the control condition.

### Electrophysiology

Neurons were incubated at DIV35 in artificial cerebrospinal fluid (ACSF) containing 125 mM NaCl, 2.5 mM KCl, 2.3 mM NaH₂PO₄, 25 mM NaHCO₃, 2 mM CaCl₂, 1.3 mM MgCl₂, 1.3 mM sodium ascorbate, 3.1 mM sodium pyruvate, and 10 mM dextrose (315 mOsm), continuously bubbled with 95% O₂ / 5% CO₂. Whole-cell patch-clamp recordings were performed using glass microelectrodes (6–10 MΩ) filled with an internal solution containing 120 mM potassium gluconate, 4 mM MgCl₂, 10 mM HEPES, 0.1 mM EGTA, 5 mM NaCl, 20 mM KCl, 4 mM ATP-Tris, 0.3 mM GTP-Tris, and 10 mM phosphocreatine (pH 7.3; 290 mOsm). Action potential firing was assessed in current-clamp mode by holding the resting membrane potential at −70 mV and applying successive depolarizing current steps of 10 pA increments with a duration of 500 ms. Voltage-gated Na⁺ and K⁺ currents were measured in voltage-clamp mode by detecting the fast inward current peak corresponding to Na⁺ currents and the delayed outward current plateau corresponding to K⁺ currents. Membrane capacitance and input resistance were calculated from current traces evoked by a 10 mV hyperpolarizing voltage step. Series resistance typically ranged between 10–20 MΩ, and recordings were excluded if series resistance exceeded 50 MΩ. Signals were recorded using Multiclamp 700B amplifiers (Molecular Devices, Sunnyvale, CA), sampled at 20 kHz, and analyzed using pClamp 10 software.

### Western blotting

Equal amounts of protein (50 µg) diluted in 20 μl of loading buffer (0.5%bromophemol blue; 10% glycerol; 10% 2-Mercaptoethanol) were loaded onto 12% SDS polyacrylamide gels (30% Acrilamide and N,N’-Methylenebisacrilamide, SIGMA). When the run was finished gels were blotted onto PVDF membranes (Whatman, USA). Membranes were blocked using 5% bovine serum albumin (BSA) in 0.05% Tween-20 in PBS (TBS-T) for 1 hour. After washing with TBS-T, membranes were incubated with the corresponding primary antibodies overnight at 4°C. Membranes were washed thrice with TBS-T and incubated with the appropriate goat secondary antibody, conjugated with horse radish peroxidase (HRP), for 2 hours at RT. Protein bands were detected using ECL (Pierce) and the Chemidoc system (Biorad). Resulting images were analyzed using ImageJ. *3D DNA-PAINT imaging and analysis.* i3Neurons were cultured on Lab-Tek II 8-well glass-bottom chambers (Nunc, 155409) at a density of 15,000 - 20,000 cells per well for 14 days *in vitro*. At DIV14, neurons were fixed using the cytoskeletal fixation protocol and subsequently immunostained for total tau and tubulin. After primary antibody incubations, samples were incubated with DNA-conjugated secondary nanobodies carrying docking strands (Massive Photonics), specific for mouse or rabbit primary antibodies. Incubations were performed for 1hr at RT at a 1:150 dilution in the manufacturer’s buffer. Samples were then incubated with a solution of 100 nm gold nanoparticles for 7 min at RT, to be used as fiducial markers for subsequent localization analysis. Imaging was performed on a custom-built widefield fluorescence microscope assembled in the laboratory of Dr. Stefani (CIBION-CONICET). The system was based on an Olympus IX-73 inverted microscope equipped with an sCMOS camera (Hamamatsu ORCA Flash 4.0). Images were acquired using a 60X objective (NA 1.45), yielding an effective pixel size of 112 nm. After selecting the imaging field and focal plane, between 30,000-60,000 frames were acquired using an imager strand complementary to the docking strand and labeled with Cy3b, diluted to 200–400pM depending on the density and frequency of blinking events. Excitation was provided by a 532nm laser at a power density of 1–2kW/cm². To improve the signal-to-noise ratio, highly inclined and laminated optical sheet (HILO) illumination was used (73). Post-processing was carried out using the Picasso software package, where single-molecule localization was performed using the maximum likelihood estimation (MLE) Integrated Gaussian method. Picasso was also used for rendering the super-resolution images. For comparative analyses, the proximal axon (defined as the AIS), distal axon, and dendrites, regions of interest were selected based on neuronal morphology, AIS length and position and projections. Briefly, an initial clustering analysis of tau localization (x, y) coordinates was performed using HDBSCAN, followed by incorporation of the z-dimension in a second clustering step using k-means. Cluster curation included a temporal filtering step, retaining only clusters whose median localization time fell within the central 66% of the acquisition window, to exclude non-specific binding and acquisition artifacts. After curation, the distance to the first nearest neighbor (d1NN), cluster area (convex hull), and number of localizations per cluster were quantified. First-nearest-neighbor distance distributions were fitted using one-dimensional Gaussian mixture models (1D-GMM).

### Statistical analysis

Statistical analyses were performed using GraphPad Prism or MATLAB. Parametric (ANOVA) or non-parametric (Kruskal-Wallis or Mann-Whitney) tests were applied appropriately following normal distribution of data, with significance depicted in asterisks as described in figure legends. Fluorescence and clustering parameters distributions were compared with Kolmogorov–Smirnov tests. For segmental velocities analyses, gaussian mixture model parameters (mode center and fraction) with their standard deviations were analyzed by comparison of the 20% non-overlapping Cis with control condition. Biological replicates are indicated as “n”.

## Supplementary figure legends

**Figure S1: hiPSC-derived glutamatergic neurons also exhibit an intrinsic AIS development pattern.** (*A*) Confocal fluorescence images of hiPSC-derived glutamatergic neurons at DIV5, 14, 25, and 37 stained against AnkG (green) and MAP2 (magenta), to visualize the AIS and the somatodendritic compartment, respectively (scale bars = 10 μm). (*B*) Quantification of the percentage of neurons with AnkG staining, (*C*) the AIS-to-soma distance, and (*D*) the AIS length. Kruskal-Wallis tests were applied (*B*: n_DIV5_=127, n_DIV14_=119, n_DIV25_=59, n_DIV37_=114; *C* and *D*: n_DIV5_=6, n_DIV14_=19, n_DIV25_=53, n_DIV37_=67, from 3 independent experiments, *p<0.05, **p<0.01, ****p<0.0001). (*E*) Quantification of the average arbitrary units of fluorescence intensity (AU) of AnkG staining at the AIS. Kolmogorov-Smirnov test was used (*E*: n_DIV5_=6, n_DIV14_=19, n_DIV25_=49, n_DIV37_=49, from 3 independent experiments, **p<0.01).

**Figure S2: Tau isoform composition does not alter basal electrical properties in human-derived neurons.** Patch-clamp recordings of DIV35 hiPSC-derived neurons infected with control, LV-PTM4R-tau or LV-PTM3R-tau. Values of (*A)* input resistance, (*B*) action potential amplitude, and (*C*) threshold obtained for each condition are shown (neurons N_control_=21, N_4R_=9, N_3R_=11, from wells from 3 independent experiments).

**Figure S3: Distribution of segmental velocities of lysosomes at the AIS.** Histograms of (*A*) anterograde and (*B*) retrograde segmental velocities of lysosomal movement at the AIS of control, LV-PTM4R- and LV-PTM3R-tau infected neurons. These distributions were fitted to a 3-Gaussian mixed model, based on the “tug-of-war/coordination” model of axonal transport.

**Figure S4: Full images of western blots used in this study.** (*A*) Mouse brain lysates, (*B*) hippocampal primary cultures, *(C)* i3Neurons timeline, *(D)* i3Neurons transduced with *transplicing* reaction for tau isoform disbalance.

**Movie S1: Lysosomal movement in hiPSC-derived glutamatergic neurons.** 30s movie (6.23 frames/sec) of a DIV14 human-derived glutamatergic neuron treated with LysoTracker dye (white) and an antibody against Neurofascin-186 (liveNF-186, green), to observe lysosomal dynamics along the AIS.

---

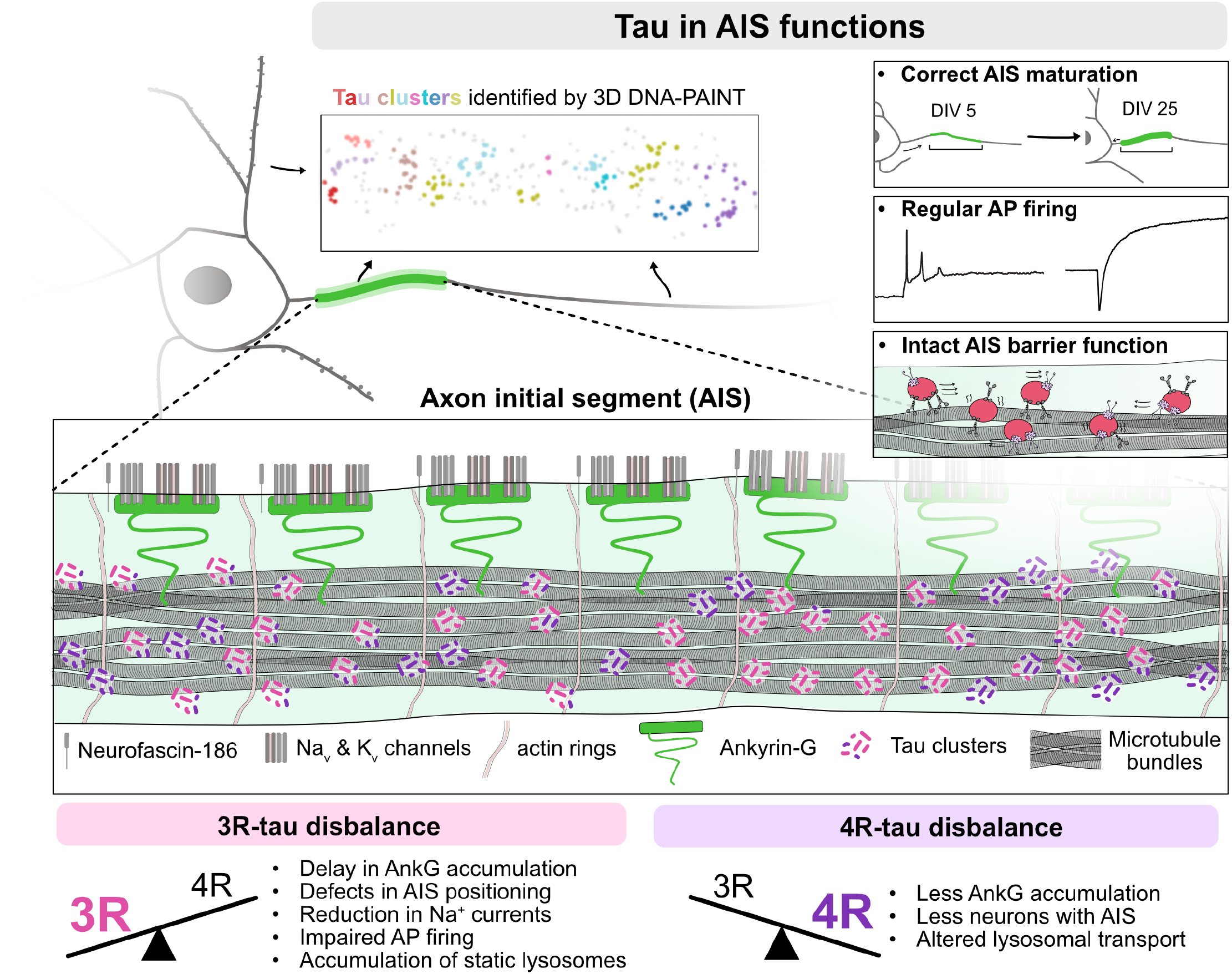

