## Supplementary figures for "Tau isoforms modulate the axon initial segment controlling axonal trafficking and neuronal excitability"

Supplementary figure 1

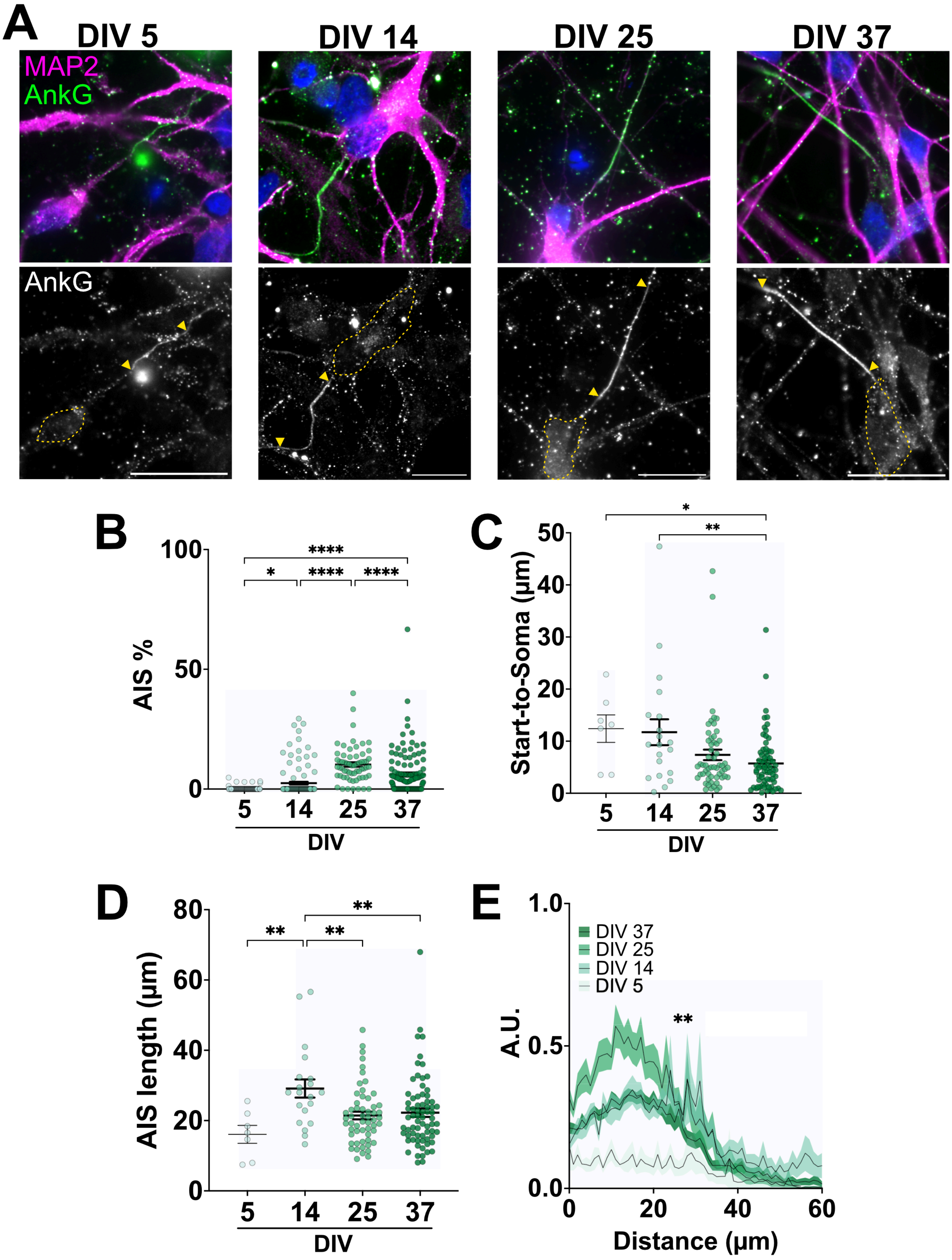

#### Supplementary figure 2

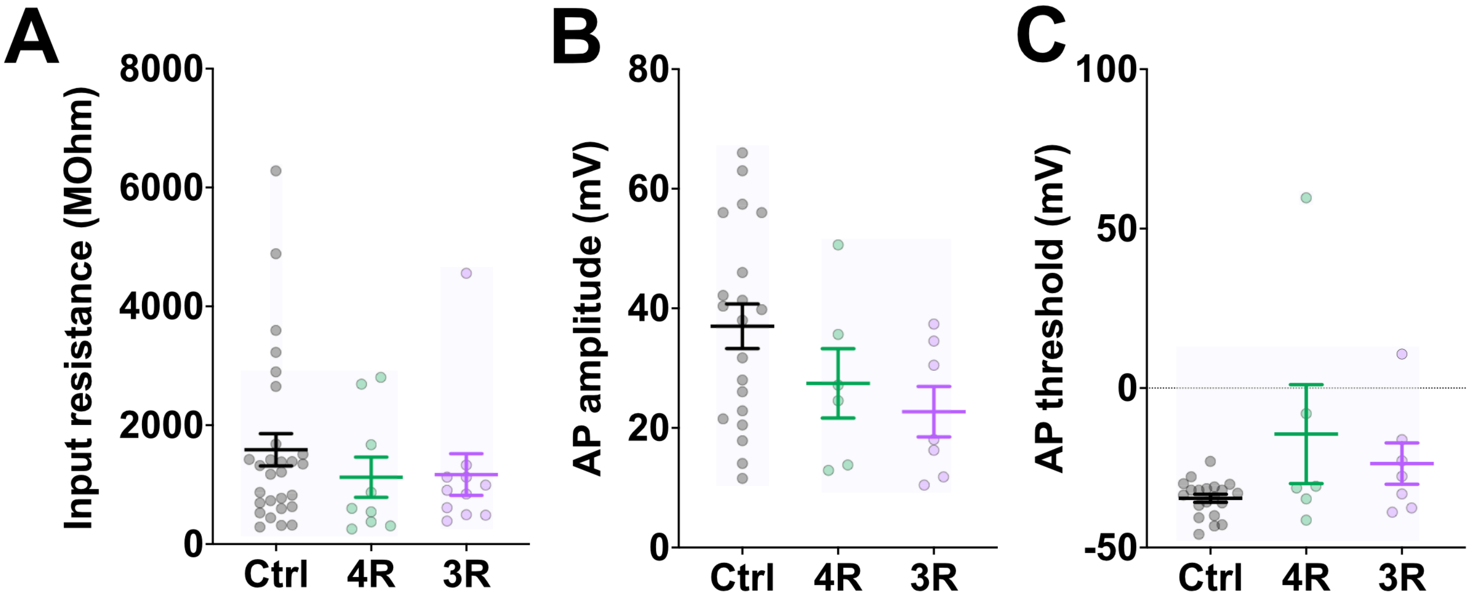

### Supplementary figure 3

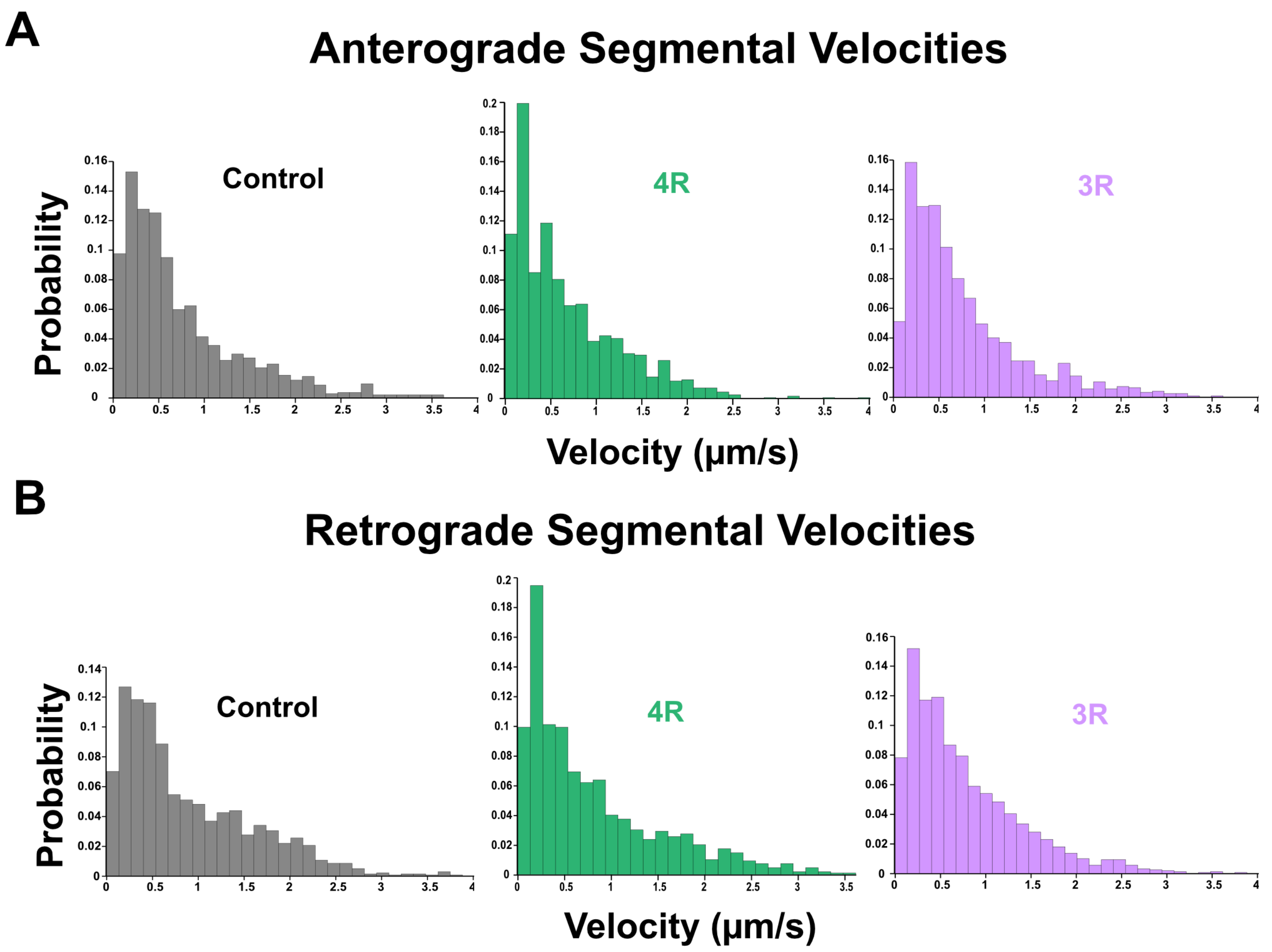

### Supplementary figure 4

#### A Brain lysates

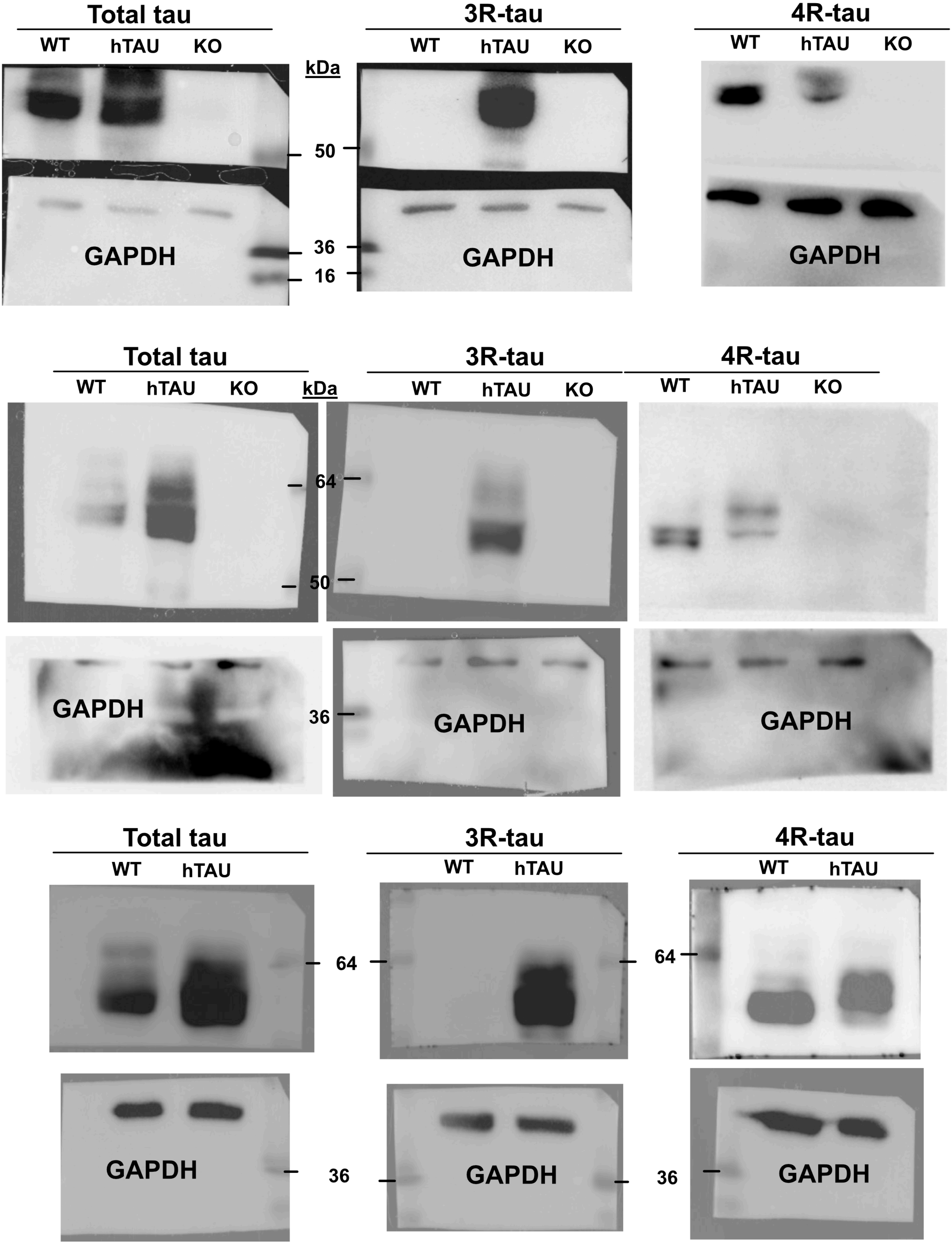

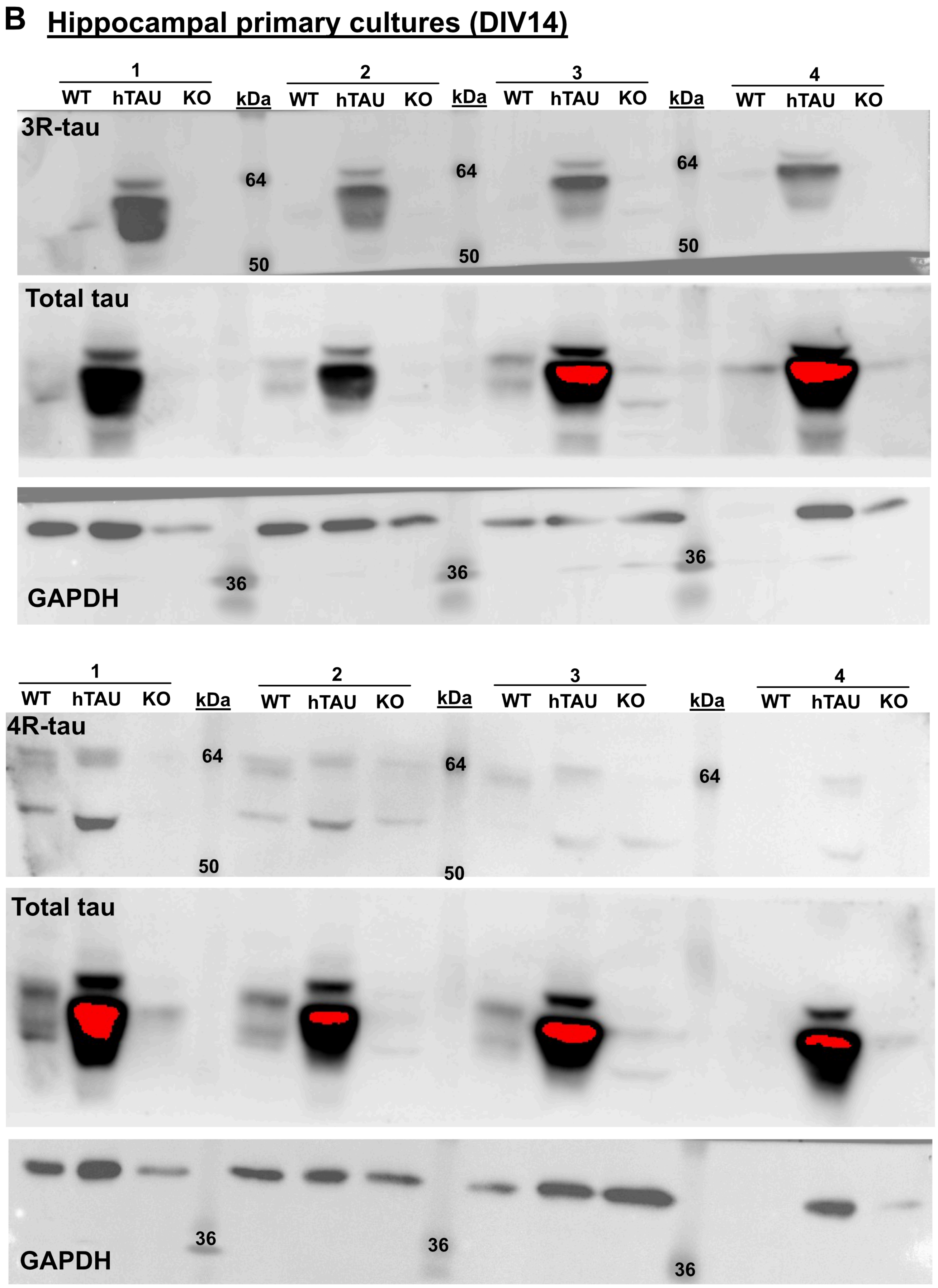

B
Hippocampal primary cultures (DIV14)
1
2
3
4
WT
hTAU
KO
kDa
3R-tau
64
50
Total tau
GAPDH
36
4R-tau

#### C i3Neurons - timeline

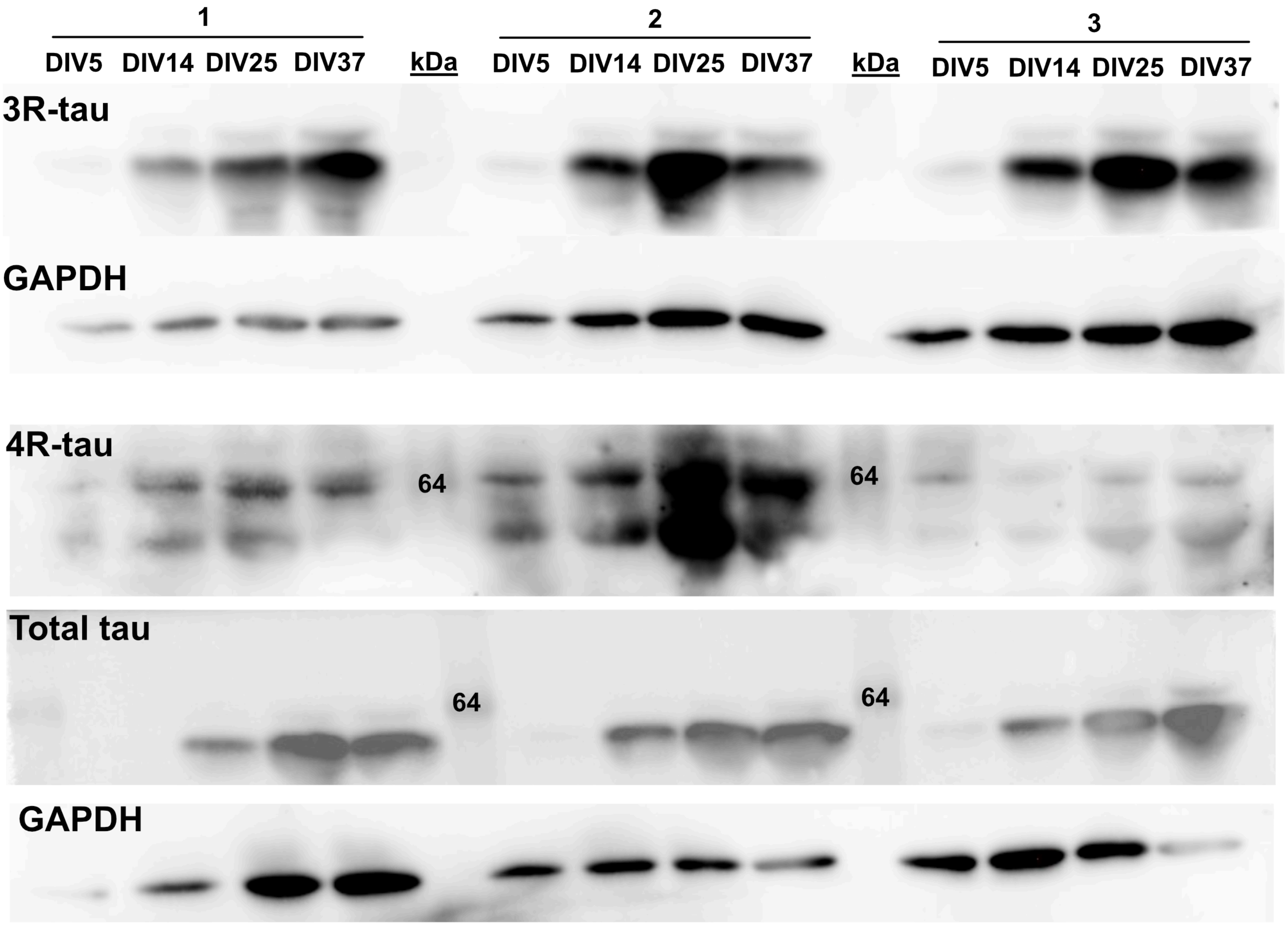

**C** i3Neurons - *transplicing* of tau isoforms (DIV14)

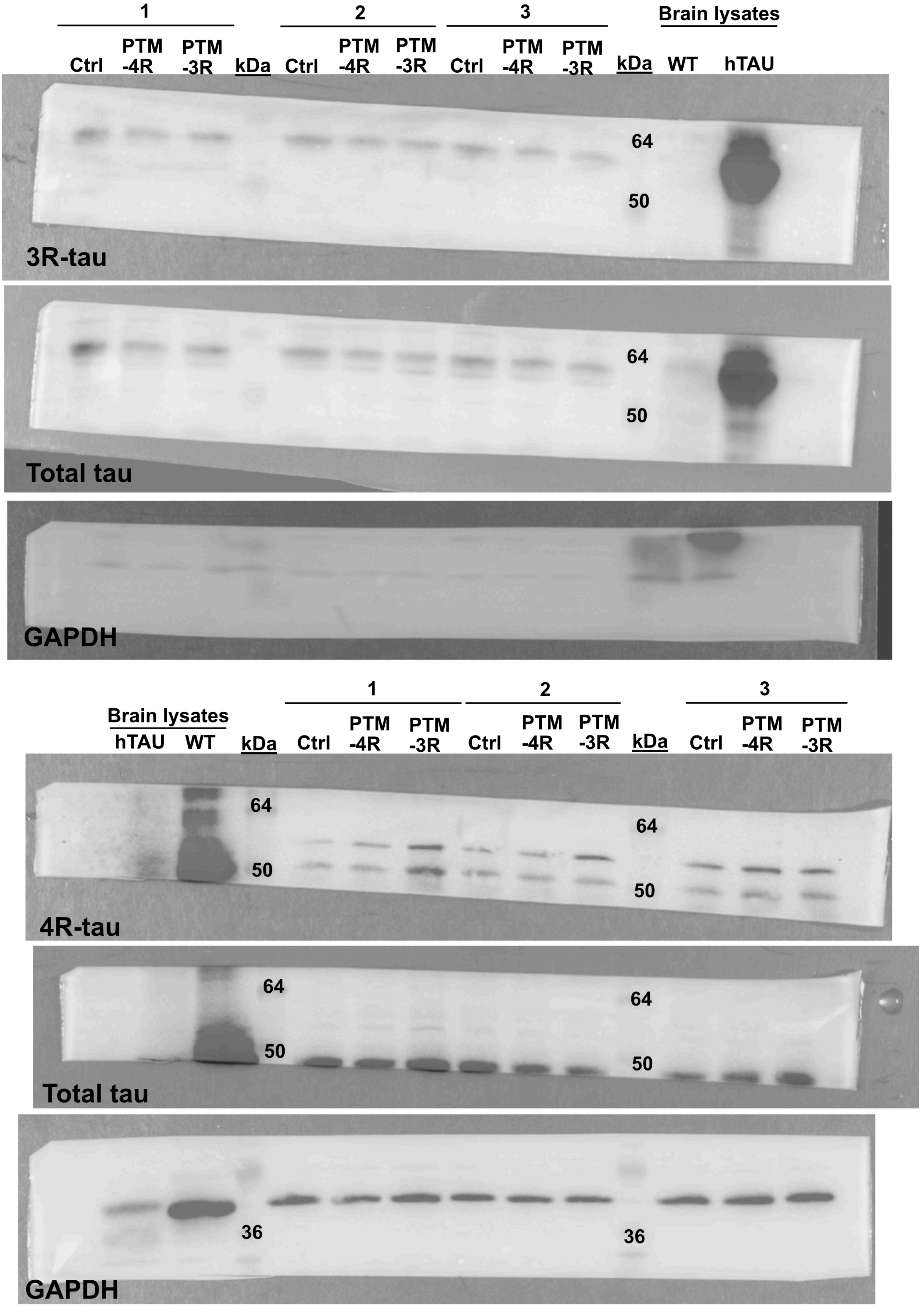

### i3Neurons - *transplicing* of tau isoforms (DIV14)

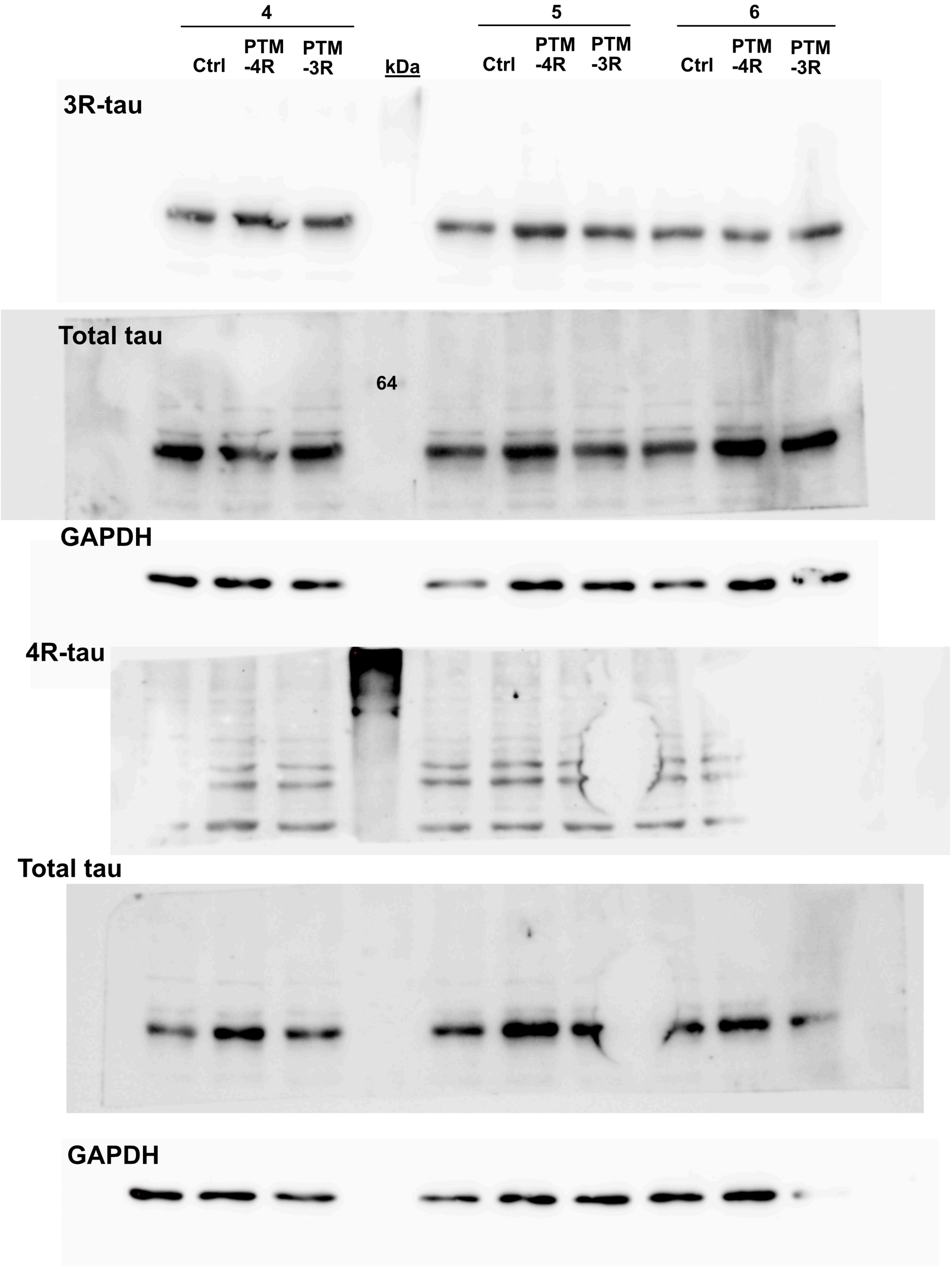
